# Dynamic brain network changes in resting-state reflect neuroplasticity: molecular and cognitive evidence

**DOI:** 10.1101/695122

**Authors:** Zhaowen Liu, Xiao Xiao, Kai Zhang, Qi Zhao, Xinyi Cao, Chunbo Li, Min Wang, Wei Lin, Jiang Qiu, Barbara J. Sahakian, Jianfeng Feng, Trevor W. Robbins, Jie Zhang

**Affiliations:** Psychiatric and Neurodevelopmental Genetics Unit, Center for Genomic Medicine, Massachusetts General Hospital, Boston, MA 02114, USA; Department of Psychiatry, Massachusetts General Hospital, Harvard Medical School, Boston, MA 02114, USA; Stanley Center for Psychiatric Research, Broad Institute of MIT and Harvard, Cambridge, MA 02138, USA; Institute of Science and Technology of Brain-inspired Intelligence, Fudan University; Key Laboratory of Computational Neuroscience and Brain Inspired Intelligence (Fudan University), Ministry of Education, PR China; Department of Computer and Information Sciences, Temple University, 1801 North Broad Street, Philadelphia PA 1912; Department of Mathematical Sciences, Fudan University, Shanghai, 200433, PR China; Shanghai Key Laboratory of Psychotic Disorders, Shanghai Mental Health Center, Shanghai Jiao Tong University School of Medicine, Shanghai, China; Center for Excellence in Brain Science and Intelligence Technology (CEBSIT), Chinese Academy of Science, Shanghai, China; Department Neurobiology, Yale Medical School, 333 Cedar St., New Haven, CT 06510, USA; Key Laboratory of Cognition and Personality (SWU), Ministry of Education, Chongqing 400715, China; School of Psychology, Southwest University (SWU), Chongqing 400715, China; Department of Psychiatry, School of Clinical Medicine, University of Cambridge, Cambridge CB2 0SZ, UK; Department of Computer Science, University of Warwick, Coventry CV4 7AL, UK; Collaborative Innovation Center for Brain Science, Fudan University, Shanghai, 200433, PR China; Shanghai Center for Mathematical Sciences, Shanghai, 200433, PR China; Department of Psychology, Behavioral and Clinical Neuroscience Institute, University of Cambridge, Cambridge CB2 3EB, UK

## Abstract

Resting-state functional brain networks demonstrate dynamic changes on the scale of seconds. However, their genetic mechanisms and profound cognitive relevance remain less explored. We identified 459 Bonferroni-corrected genes, by associating temporal variability of regional functional connectivity patterns with Allen Brain gene expression profiles across the whole brain. These genes are partially verified in developing human brain gene expression in the BrainSpan Atlas, and are found to be involved in the enrichment of short- and long-term plasticity processes. The former process depends on synaptic plasticity, involving ion transmembrane transport, action potential propagation, and modulation. The latter process depends on structural plasticity, including axonal genesis, development, and guidance. Results from a longitudinal cognitive training study further revealed that baseline variability of the hippocampal network predicted cognitive ability changes after three months of training. Our genetic association results suggest that the short-term plasticity processes may account for the rapid changes of regional functional connectivity, while the underlying long-term plasticity processes explain why temporal variability can predict long-term learning outcomes. To our knowledge, this is the first demonstration that measuring the dynamic brain network can lead to a non-invasive quantification of neuroplasticity in humans.

## Introduction

Over the last decade, researchers have tried to characterize and understand the dynamic changes of functional brain networks in both resting and task-related fMRI to shed new light on the dynamics of spatiotemporal organization of spontaneous or evoked brain activity ^1, 2, 3, 4, 5, 6, 7^. The dynamic changes of brain networks are related to cognition ^8, 9, 10, 11, 12, 13, 14, 15^, neuromodulatory systems^16, 17, 18, 19^, consciousness ^20,21^ and are altered in various psychiatric disorders ^22, 23, 24, 25^.

Whilst a considerable amount of work has been devoted to characterizing dynamic functional connectivity (FC) or network changes in spontaneous brain activity and connectivity, little attention has been paid to the underlying genetic mechanisms ^26^.The most recent efforts to integrate transcriptomic data from the Allen Human Brain Atlas into fMRI studies focused on understanding the molecular basis of static functional brain organization or structure ^27^. For example, gene expression profiles associated with regional BOLD or electrophysiological activity ^28, 29^, functionally connected regions ^30^, hubs ^31^, networks ^32, 33^, cortical thickness and myelination ^34^, and fiber connectivity ^35^ have all been characterized, revealing specific genetic signatures over different levels of functional organization of the brain. In addition, the behavioral significance of the spontaneous fluctuation of the brain network remains unresolved. For instance, it is not clear how baseline level of dynamic changes of functional brain networks may confer adaptability and plasticity for future learning.

In this work we aimed to understand both the biological basis and behavioral implications of dynamic changes of the brain network in resting-state fMRI. First, we explored the molecular basis of dynamic changes of functional brain networks by associating the topography of temporal variability, a recently proposed dynamic brain network metric ^36^, with whole-brain transcriptomic profiles provided by the Allen Brain Atlas data. We identified more than 400 genes whose expression in the brain correlated significantly with topography of temporal variability, and these genes are found to be enriched in biological processes relevant to short-term and long-term plasticity.

We further investigated the cognitive implications of spontaneous, dynamic brain changes in predicting future learning outcomes, using longitudinal cognitive training data from an aged cohort. We found that the dynamic network changes of hippocampus at baseline can predict changes in a visuospatial/constructional behavioral index after 3 months of cognitive training, suggesting that the dynamic metric probably reflects long-term plasticity or learning ability. At the molecular level, the biological pathways identified from genes, that are highly correlated with dynamic brain changes, explain why spontaneous brain network show temporal variability in resting-state and how these changes can predict long-term learning outcomes. Overall, we explored dynamic brain network changes from genetic, neuroimaging and cognitive perspectives, and our results suggest that dynamic brain changes in resting-state can measure both short- and long-term neuroplasticity.

## Results

### Topography of dynamic functional connectivity changes

We used a metric called temporal variability to quantify the dynamic changes in functional connectivity associated with a given brain region ^36^. Greater temporal variability of a region indicates that its functional connectivity with other areas is changing frequently ^36^. The topography of temporal variability for the whole brain was obtained using HCP data and HCP_MMP1.0 multi-modal parcellation.^37^ HCP_MMP1.0 parcellation provides an invaluable neuroanatomical framework for 360 areas in the group average and in individual subjects; therefore, it is more refined and reliable.

At the group level, the orbitofrontal complex (OFC) and posterior OFC (pOFC), piriform olfactory cortex, insular cortex, cingulate (Subgenual and anterior cingulate cortex/MPFC), the inferior medial temporal region (including the hippocampus/parahippocampus, presubiculum (PreS), entorhinal cortex (EC), peri-entorhinal and ectorhinal complex (PeEc), as well as temporal pole) had the highest temporal variability (see Fig. 1 and Supplemental Table 1 for details, and Supplemental Table 7 for the names of the regions and abbreviations). In contrast, the early visual cortex (V2, V3 and V5 V6, lateral occipital cortex), premotor and sensory motor cortex, as well as inferior and superior parietal cortex, including intra-parietal sulcus, had lower temporal variability.

**Figure 1.**
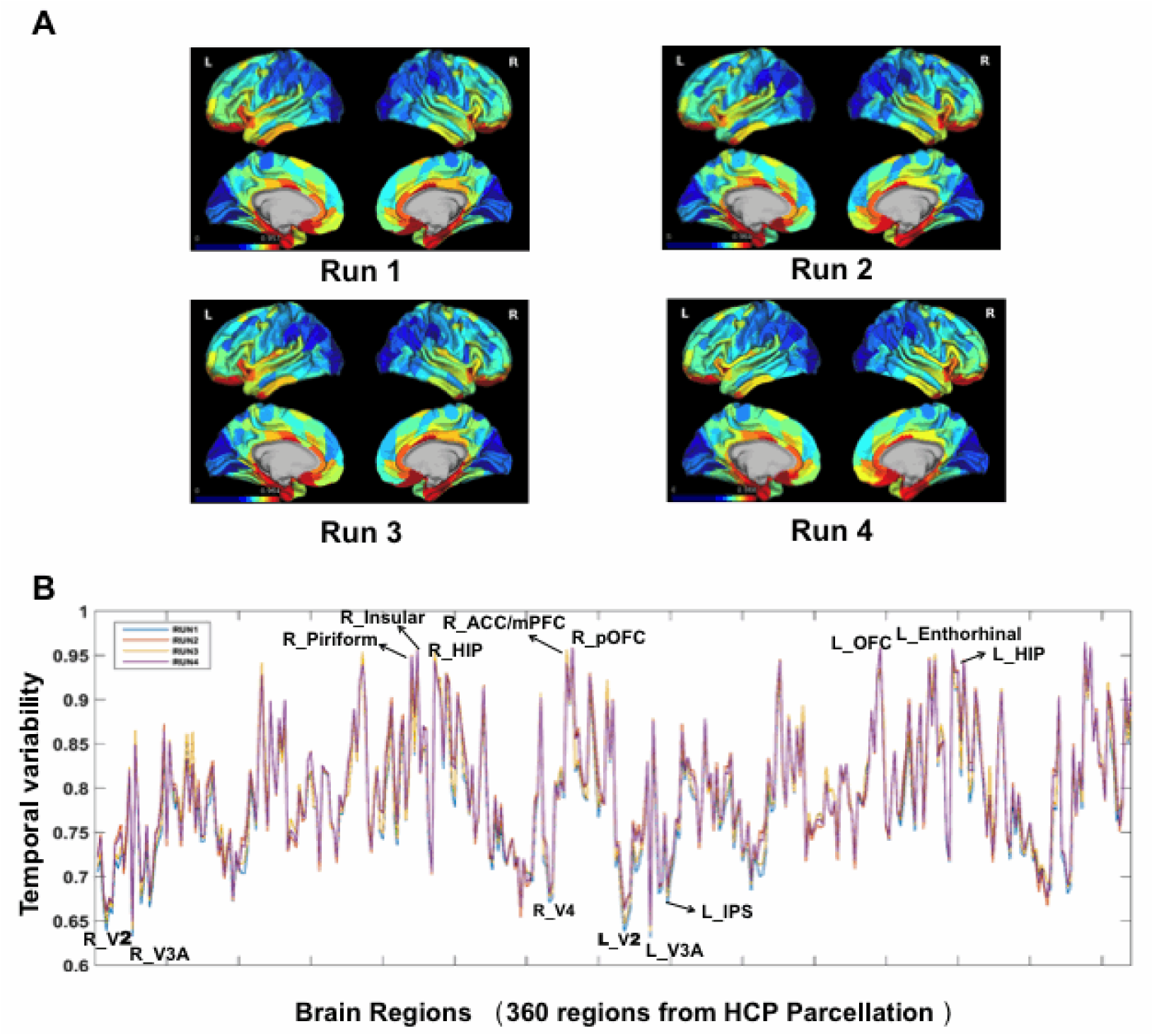
Whole-brain topography of temporal variability obtained from HCP data. The brain map is averaged across all 864 subjects and for each of the 4 scans. A. The temporal variability demonstrated high similarity across 4 scans for 360 brain regions. B. Regions with highest (like hippocampus, orbital frontal gyrus and ACC/MPFC) and lowest (like visual areas) temporal variability are shown.

Importantly, we found that the medial prefrontal cortex (PFC) demonstrates higher variability than the dorsal PFC. Ventral and medial PFC have reciprocal projections with olfactory circuitry, insular cortex and limbic brain areas, all having high temporal variability, while the lateral PFC interconnects with visual, auditory and somatosensory association cortices with low temporal variability. The medial or ventral PFC represents the internal world and is active in the resting-state, thus potentially demonstrating high variability. The lateral PFC, in contrast, represents the external world. Although the lateral PFC has less variability compared to the medial PFC in resting-state, we hypothesize that it will show higher variability in a task-realted state owing to its flexible hub role ^7^.

High variability in association cortex and limbic systems and low variability in unimodal sensory motor and visual areas were replicated in different scans in HCP data (Figure 1B), and also in UK Biobank datasets (Supplementary Table 2). The temporal variability demonstrates a high correlation between HCP data and UK Biobank data (>0.9, see Supplemental Table 2), across all brain regions (using AAL2 template). These results indicate the reliability of temporal variability as a measure of dynamic FC changes.

### Genes significantly correlated with dynamic functional connectivity changes of the brain

Temporal variability quantifies the dynamic changes of the functional connectivity profile associated with a given brain region, allowing for convenient association with whole-brain gene expression data. We correlated whole-brain topography of temporal variability with whole-brain transcriptomic profiles for about 20738 genes (Fig. 2) and found 324 genes, the expression of which in the brain showed significant negative correlation, while 135 genes showed positive correlation (Bonferroni correction; see Fig. 3A and Supplemental Table 3). Gene Ontology (GO) analysis showed that negatively related genes are functionally enriched in 48 biological processes (Bonferroni correction) related to synaptic plasticity. These genes are mainly involved in cell signaling, such as ion transport (15 terms, including various cation, e.g., potassium and sodium, and anion transport), action potential and its propagation (5 terms), peptide and hormone secretion, and transport and regulation (12 terms) (see Fig. 3A and Supplemental Table 4). The most significant term was multicellular organismal signaling (p=1.82e-8). Meanwhile, the biological processes in which the positively related genes are enriched processes that are probably more related to structural plasticity including axonal development, guidance, axonal- and neurogenesis, and neuron differentiation (p<0.001, uncorrected) (see Fig. 3A and Supplemental Table 5), with the most significant biological processes being cGMP-mediated signaling (p=5.8E-5) which plays a role in regulating synaptic plasticity ^38^.

**Figure 2.**
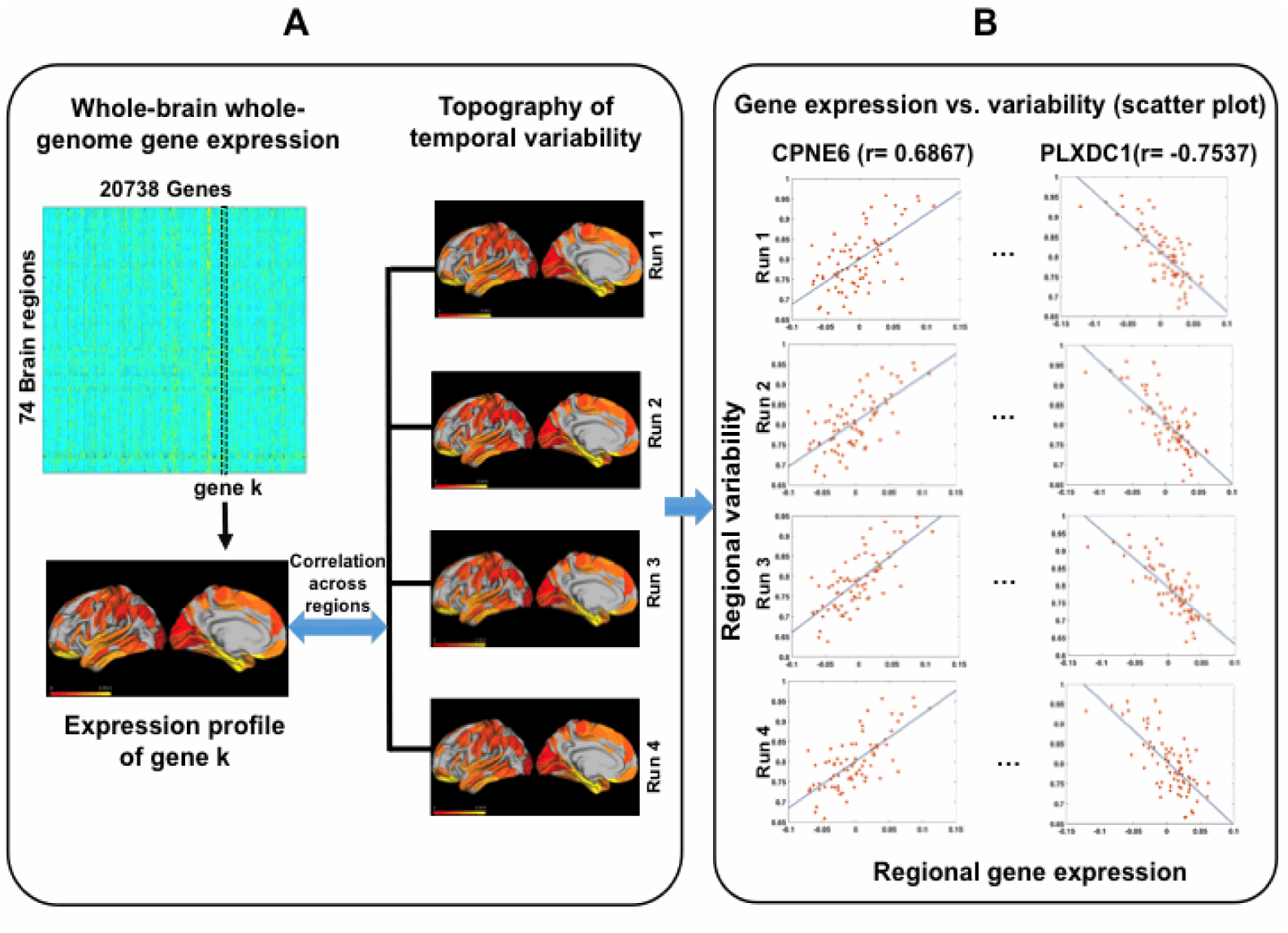
Schematic illustration of correlation analysis between whole-brain temporal variability from resting-fMRI and whole-brain gene expression profile. A. Correlation analysis between whole-brain topography of temporal variability and whole-genome gene expression. For each gene, e.g., the *k*th gene, we calculated the correlation coefficient between its expression in all brain regions and the corresponding temporal variability. Both expression and temporal variability in each brain region were averaged over the whole group of subjects. The correlation analysis was performed for each of the 4 fMRI scans, and those genes that passed the Bonferroni correction in each scan were kept for further analysis. B. Two genes, the expressions of which were most correlated with temporal variability, were shown with the corresponding scatter plots. The most positively correlated gene is CPNE6 (r=0.6867), and the most negatively correlated gene is PLXDC1 (r=-0.7537).

**Figure 3.**
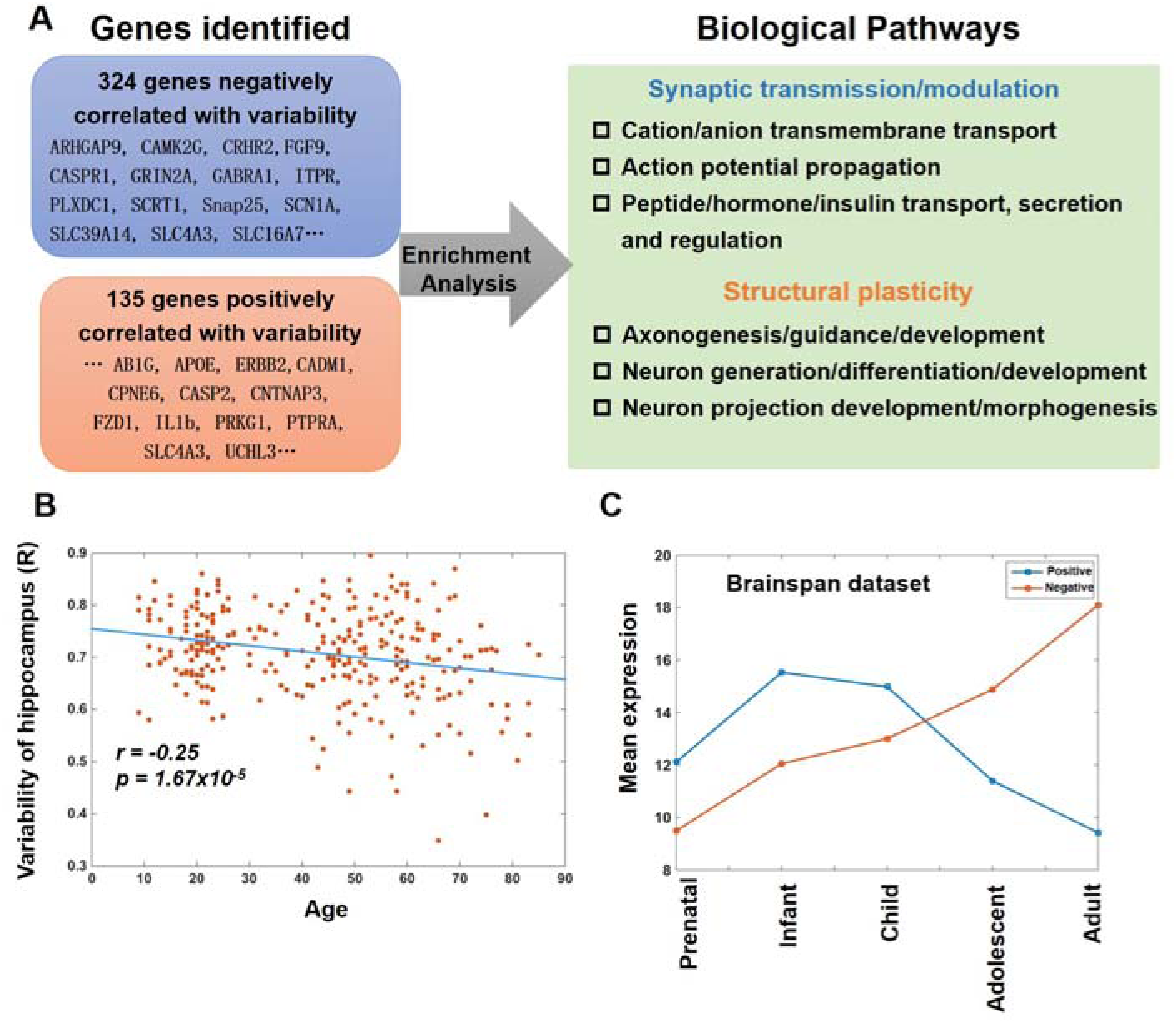
Biological pathways for genes significantly correlated with temporal variability and changes of expression of these genes with age. A. Biological pathways for genes positively and negatively correlated with temporal variability of functional brain networks. Some representative genes are also listed in the left column, and the full gene lists can be found in Supplemental table 3. B. Changes of temporal variability with age for right hippocampus. Window lengths were chosen from 20-30s in calculating the temporal variability from fMRI data. C. Changes of expression of genes over different stages of human development in BrainSpan data. The genes identified in Allen Brain Atalas were divided into two groups: negatively/positively related to temporal variability. The mean expression of genes in the two groups was shown. As can be seen in B and C, both variability and genes positively related to variability show the same decreasing trend with age, which verifies that the genes we identified in Allen Brain Atalas are also correlated with variability in BrainSpan dataset.

We found that a notable number of genes, the expression of which is correlated with dynamic network changes, were implicated in neuroplasticity and learning (see list in Fig. 3A). For example, for negatively related genes, the most significant gene was PLXDC1, which encodes PLXDC1 transmembrane proteins that act as cell-surface receptors for Pigment Epithelium Derived Factor (PEDF), a neurotrophic factor ^39^. ITPR1, which binds with DISC1 and regulates its dendritic transport ^39^, SNAP25, is a key component of the synaptic-vesicle fusion machinery ^40^, and ARHGAP9, which controls synapse development, all play a role in synaptic plasticity. GRIN2A and GABRA1, which are NMDA and GABA receptors, respectively, are key mediators of plasticity, and Caspr1 regulates the content of AMPA receptors at synapses ^41^. Other genes, such as SCN1A, which provides instructions for making sodium channels; SLC16A7, a postsynaptic protein involved in long-term memory; SLC4A3, a membrane transport protein and CAMK2G, are crucial for plasticity at glutamatergic synapses.

The gene most positively correlated with temporal variability was CPNE6, which has been reported to play a role in synaptic plasticity ^42^. Other positively related genes included APOE, which affects neurogenesis and synaptic plasticity ^43^ and FZD1 that regulates adult hippocampal neurogenesis ^44^. Also, CNTNAP3 regulates synaptic development ^45^, UCHL3 is implicated in the control of cell cycle/growth and synaptogenesis, and PTPRA is involved in numerous neurodevelopmental processes. Finally, ERBB2 is involved in Nrg/Erbb signaling networks regulating the assembly of neural circuitry, myelination and synaptic plasticity ^46^.

### Correlation between variability-related gene expression and age

To partly verify the identified genes whose expression was significantly correlated with temporal variability in the Allen Brain Atlas dataset, we also analyzed the change of these genes’ expression with age using the Brain Span human development data. First, we found that the temporal variability of most brain regions demonstrated significant negative correlation with age (Supplemental Table 6), most prominently in the hippocampus (Fig. 3B). Second, for the genes identified in the Allen Brain Atlas dataset that are positively or negatively related to variability, we calculated their mean expression each age category respectively within the BrainSpan dataset. We found that gene expression that was positively related to dynamic functional connectivity changes significantly decreased with age. In contrast, whereas gene expression that was negatively related to dynamic functional connectivity changes, increased from infancy through to adulthood (see Fig. 3C for details).

These findings suggested that those genes in the Allen Brain Atlas dataset that were identified to be correlated with temporal variability were also correlated significantly with variability in the BrainSpan data related to similar age-related changes, which partly confirms the genes found in Allen Brain Atlas dataset. As plasticity reduces with aging ^47^, these results also confirm the close relationship between dynamic functional connectivity changes, neuroplasticity and plasticity-related genes.

### Short-term dynamic FC changes predict long-term learning outcomes

We tested whether current brain network variability is predictive of future learning outcomes in order to understand the cognitive significance of the temporal variability of functional connectivity changes. To determine this, we measured changes in cognitive scores following a period of training. The study assessed an aging cohort in a longitudinal cognitive training study ^48, 49^. Our study included 26 subjects from the community (70.38□±□3.30 yrs) who underwent training of multiple cognitive domains including memory, reasoning, problem-solving strategies, and visuospatial map-reading skills, for 1 hour twice a week for 12 weeks.

We measured the baseline level of temporal variability from resting-fMRI for each brain region and correlated this with the change in cognitive score after 12 weeks of training, as measured by the Repeatable Battery for the Assessment of Neuropsychological Status score (RBANS, Form A).^50^ The RBANS is a widely used test for cognitive assessment. It includes a visuospatial/constructional index composed of a complex figure copy and judgment-of-line orientation task, reflecting a central cognitive ability. Of all 90 brain regions, only the baseline level variability of the right hippocampus showed a significant positive correlation with the change in visuospatial/constructional index (r=0.47, p= 0.01, FDR corrected).

## Discussion

The dynamic changes of functional brain networks in spontaneous resting-fMRI have become increasingly recognized as having important functional implications for cognitive performance, development, aging, and disease ^51^. Despite the growing body of work and interest in characterizing such dynamic changes, the underlying genetic mechanisms have to date been poorly addressed. Our genetic association analysis revealed neuroplasticity-related biological processes underlying dynamic functional connectivity changes. Furthermore, our longitudinal cognitive training data revealed that baseline temporal variability could predict learning outcomes within a window of several months for an aging cohort. We explored dynamic brain network changes from genetic, neuroimaging and cognitive perspectives, and together our results indicated that temporal variability measured from resting-state brain networks can serve as potential endophenotypes that reflect neuroplasticity.

Dynamic FC changes reflect a fast plasticity process. Conceptually, dynamic changes of functional connectivity patterns on the scale of seconds are consistent with a new form of very rapid plasticity, termed Dynamic Network Connectivity (DNC) ^52, 53, 54^. Although the precise mechanisms of dynamic brain network changes remain unclear, it is believed that brain network reconfiguration can be rapid and transient and related to cognitive ability or task performance ^51^. At the molecular level, a recent work found consistency between the “dynamics” of functional connectivity calculated using calcium and hemodynamic signals, suggesting a neuronal origin of the temporal fluctuations of hemodynamic functional connectivity ^55^. In this process, the strength of neural-network connections can be rapidly increased/decreased over a short time scale, as manifested by changes in coherence synchrony and phase-locking membrane potentials in remote neuronal ensembles ^54^, which is highly consistent with our variability measure of functional connectivity in resting-fMRI.

The enrichment analysis of genes closely linked to dynamic connectivity changes provides a molecular explanation for the fast changes in resting-state functional connectivity. Biological processes, including synaptic transmission and modulation (see Fig. 3A and Fig. 4 A, C), are essential for maintaining and modulating functional connectivity. The neural network connectivity and its changes are closely related to these synaptic transmission processes that rely upon the availability of neurotransmitter transporters in presynaptic axonal terminals, the release of neurotransmitters, the activation of postsynaptic receptors and ion channels, and even the longer term changes of signaling cascades such as calcium-cAMP signaling, as well as modulation by various neuropeptides and hormones.

**Figure 4.**
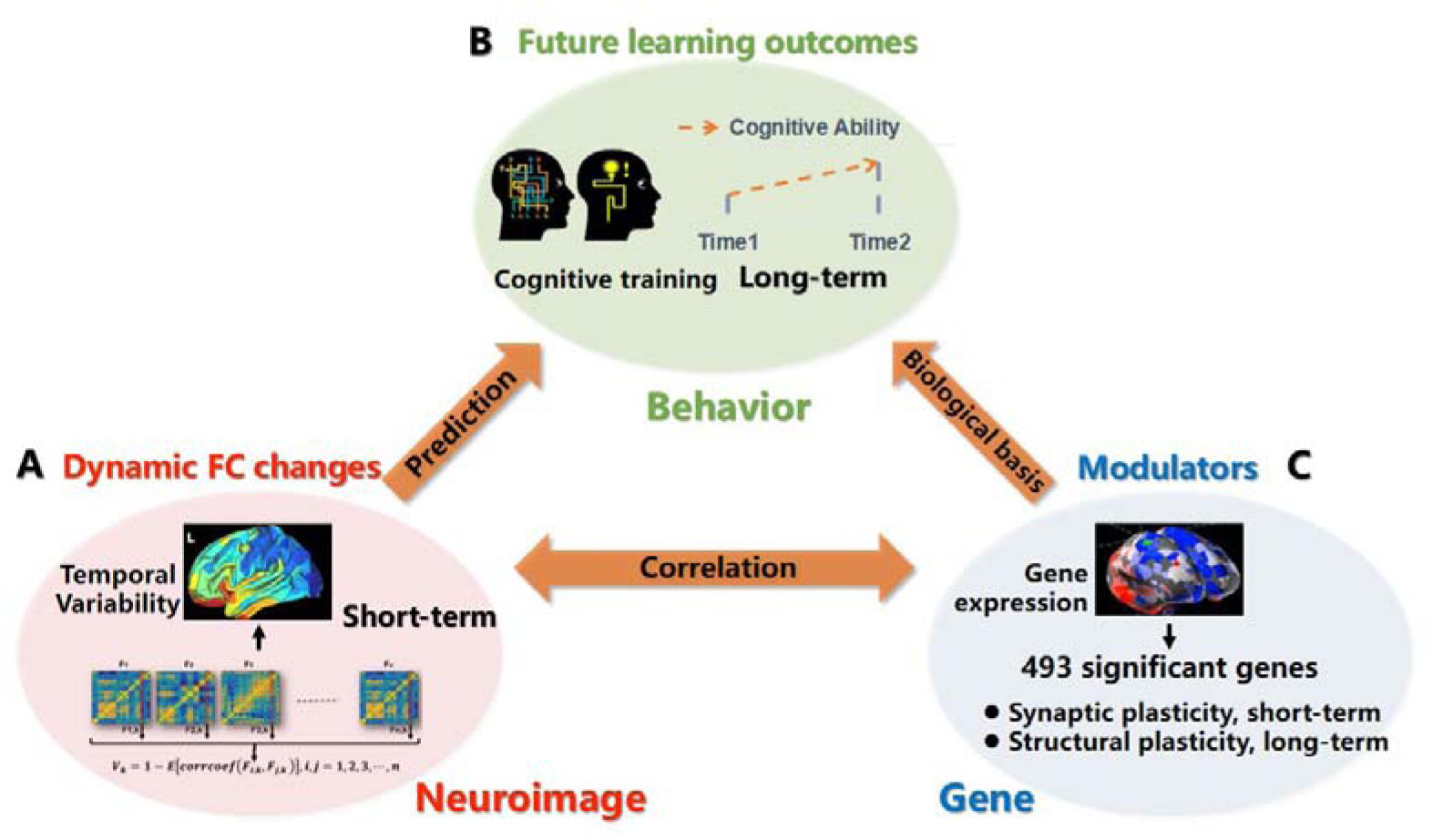
Schematic illustration of the relationship among dynamic brain network changes, i.e., temporal variability (A), underlying genes and biological pathways (B), and long-term learning outcomes (C). By correlating temporal variability of a brain region (A) with gene expression (B) across the whole brain, we obtained 493 significant genes (Bonferroni correction) enriched in short- and long-term plasticity, i.e., synaptic plasticity and structural plasticity (C). Dynamic brain network measures of right hippocampus (A) can also predict future learning outcomes, i.e., changes in visuospatial/constructional index after 3 months of learning (B). The genetic basis for temporal variability involved both short- and long-term plasticity (C), which explains why short-term, spontaneous functional brain networks changes can predict long-term learning outcomes.

Dynamic FC changes are also intimately related to long-term plasticity processes, as we found that the baseline variability relative to functional connectivity of the hippocampus can predict changes in cognitive performance after 3 months of training for an aging cohort (see Fig. 4A and B). Learning and memory occur through modifications in the strength of neural circuits, or neuroplasticity, particularly in the hippocampus. Therefore, our results suggested that the variability of FC changes in resting-fMRI may reflect neuroplasticity at the behavioral level. Whilst a number of studies have demonstrated the behavioral and cognitive relevance of dynamic changes in functional connectivity ^51^, it should be noted that these studies showed spontaneous functional connectivity dynamics. For example, their studies reported the impact of neural network structure on concurrent behavior or how pre-stimulus functional connectivity is related to immediate task performance ^8, 51, 56, 57^. Few studies have shown the capability of dynamic network endophenotypes to predict future learning, especially after a long period of cognitive training.

Our findings explain why spontaneous, short-term changes of current resting-state networks are indicative of the capability to learn in the future, or long-term plasticity. Dynamic changes in functional connectivity, or “spontaneous fluctuations in intrinsic connectivity networks”, have been postulated to optimize the brain’s readiness to respond to similar inputs in the future ^58^. Furthermore, these “intrinsic coupling modes” that change dynamically over time are hypothesized to represent spatiotemporal coupling patterns that adapt through use-dependent plasticity ^59^. In this regard, we believe that large FC variability in resting-state indicates patterns of functional connectivity that could potentially accommodate various cognitive demands. Therefore, large variability facilitates preparation of functional brain networks for participation in processes involved in future learning.

As evidenced in our longitudinal cognitive training data, our gene association analysis additionally provides molecular evidence that explains why dynamic FC changes are predictive of long-term learning outcomes. We found that genes closely related to dynamic FC changes are mainly enriched in two forms of neuroplasticity process (see Fig. 3A). The first appears to be related to synaptic plasticity or short-term plasticity, for example, alterations in existing synapses, including synaptic transmission, neurotransmitters, receptors, and modulators. (Fig. 3A upper panel). As discussed above, these processes are related to fast changes in functional connectivity. The second is probably related to structural plasticity or long-term plasticity, involving structural changes, such as the formation and development of postsynaptic assembly (dendritic remodeling), presynaptic assembly (axonogenesis, sprouting, and pruning), and soma (neurogenesis) (see Fig. 3A, lower panel). These structural changes, such as establishing a connection between spines and synapses, are correlated with the consolidation or maintenance of learning and memory ^60^. Spine formation and synaptogenesis are triggered by cell surface adhesion molecules (CAMs) and their component proteins ^61^. Consistently, we have identified most CAMs and proteins in our gene association analysis, e.g., ERBB2, CADM1, and VANGL (see Fig. 3A and Supplemental Table 3).

In summary, we explored dynamic functional brain network changes from genetic, neuroimaging and cognitive perspectives, and our results suggested that the biological processes underlying spontaneous dynamic FC changes in resting-fMRI involves both short- and long-term plasticity processes. This explains why fast-scale dynamic FC changes can predict long-term learning process at the molecular level. The temporal variability metric from resting-fMRI can be used in future to quantify the level of neural plasticity and learning in a range of age groups and environmental contexts. Our findings have important clinical implications in regard to early changes in plasticity in both healthy young people and the elderly, and in neurodevelopmental disorders and neurodegenerative diseases.

## Methods

### fMRI data from HCP

For our analyses, we used data from the 900 Subjects Release (S900) of the Human Connectome Project (HCP). The S900 collected 897 healthy adults from August 2012 to Spring 2015. Details can be found at https://www.humanconnectome.org/study/hcp-young-adult/. Using multi-modal imaging data from the HCP and a machine learning method, we identified a multi-modal parcellation (MMP) consisting of 180 regions for each hemisphere ^37^. We used the 210v (210-subject validation) group MMP parcellation (https://balsa.wustl.edu/88mp) in our analyses.

The 3T functional magnetic resonance imaging (rfMRI) data were acquired with a spatial resolution of 2 × 2 × 2 mm and temporal resolution of 0.7 s. The detailed acquisition protocol used for rfMRI data was comprehensively described in ^62^. We used all four runs of fMRI data collected over the course of two sessions. For each session, the phase encoding of oblique axial acquisitions was obtained in a right-to-left (RL) direction in one run and in a left-to-right (LR) direction in the other run. The number of subjects collected for each run varied from 828 to 870. For all sessions, all the data from both phase-encoding runs were used to calculate temporal variability.

### fMRI data for longitudinal cognitive training

For cognitive training fMRI data, 26 healthy older adults with normal functional capacity, living independently in the community were recruited to a longitudinal cognitive training study ^49^ between March 2008 and April 2008 in Tongji Hospital. These subjects aged 65–75 years, with education ≥ 1 year, had no hearing, vision, or communication problems and no severe physical, neurological, or psychological disease or obvious cognitive decline, such as brain cancer, major depressive disorder, schizophrenia, and AD. Participants completed a one-hour multi-domain cognitive training twice a week for 12 weeks, targeting memory, reasoning, problem-solving strategies, and visuospatial map-reading skills. All participants were initially given a cognitive capacity assessment (baseline). Three months after intervention, another cognitive assessment was conducted. The measurements included the Repeatable Battery for the Assessment of Neuropsychological Status (RBANS, Form A) ^50, 63^, which has been shown to have good reliability and validity in Chinese community-living older people ^63^.

Scanning was performed using a Siemens 3.0 Tesla Allegra scanner (Erlangen, Germany) at East China Normal University, Shanghai, China, both at baseline and after 3 months of training. Functional images were acquired using a single-shot, gradient-recalled echo planar imaging sequence (repetition time = 2000 ms, echo time = 25 ms and flip angle = 90°). Thirty-two transverse slices (field of view = 240 × 240 mm^2^, in-plane matrix = 64 × 64, slice thickness = 5 mm, voxel size = 3.75 × 3.75 × 5 mm^3^). Subjects were instructed to rest with their eyes closed and not to fall asleep. A total of 155 volumes were acquired, resulting in a total scan time of 310 s.

### fMRI data for exploring variability changes with age

In order to evaluate how temporal variability of brain networks changes with age, we used resting-fMRI data from http://fcon_1000.projects.nitrc.org/indi/pro/nki.html, which consisted of individuals between the ages of 4 and 85 years old. After data quality control, 298 subjects were left with a mean age of 42.75 y with a standard deviation of 19.57 y.

### fMRI data preprocessing

For the HCP dataset, the resting-state fMRIs were run through minimal preprocessing pipelines ^64^. Next, the data were resampled to the standard grayordinates space according to the areal-feature-based MSM surface registration. Subsequently, the ICA+ FIX approach was performed for each volumetric time series to remove the spatially specific temporal artefacts. The approach includes the following procedures: 1) linear trend removal, 2) MELODIC-independent component analysis (ICA), 3) FMRIB’s ICA-based Xnoiseifier (FIX) to separate noise from signal, 4) regression out of the data noise and the 24 motion parameters. Here we focus on the 64984 grayordinates that belong to the cerebral cortex. Three hundred sixty regional time series were extracted by averaging voxel time series within each of the MMP regions, and the names of the regions and their corresponding abbreviations are listed in Supplemental Table 7.

### Temporal variability: measure of dynamic FC changes of a brain region

We used a dynamic neuroimaging endophenotype called temporal variability that we previously proposed ^36^ for association with the brain’s gene expression. The temporal variability of a brain region characterizes the dynamic changes of the functional connectivity patterns of a given brain region across different time windows ^36, 65^. To obtain the temporal variability, we first segmented all BOLD signals into non-overlapping windows (length l). The whole-brain functional network F_i_ in the *i*th time window was then constructed, using Pearson correlation as the measure of FC. The functional architecture of region *k* at window time *i* is denoted by *F_i,k_*, which represents all the functional connections of region *k*. The variability of an ROI k is defined as

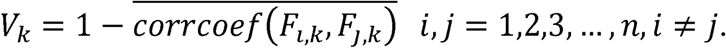

We calculated *V*_k_ at a number of different window lengths (l=equal to 25, 26, 27…50 seconds) and then took the average value as the final variability. These window lengths were chosen because window sizes around 30–60 s lead to robust results for cognitive states ^66^ and topological descriptions of brain networks ^67^. It should be noted that variability at different window length is highly correlated, indicating that the choice of window length is not crucial.

Variability characterizes the flexibility of the regional functional architecture. It reflects the region’s ability to reconfigure itself into different functional communities, or the flexibility in the functional integration/coordination with different neural systems. The larger the temporal variability of an ROI, the more different functional communities it will be involved in at different times.

### Adult brain gene expression data and preprocessing

We used the gene expression data of six human brains from the Allen Human Brain Atlas (http://human.brain-map.org) for association analysis with the dynamic, neuroimaging endophenotype. We used an improved normalization process implemented in March 2013. Two of the brains had both hemispheres, and two had only the left hemisphere. In total, nearly 4000 unique anatomic samples with expression profiles of 20738 genes were obtained. The details of microarray information and data normalization are available at http://help.brain-map.org/display/humanbrain/documentation/.

For each AHBA issue sample, we create a 6mm sphere ROI in the MNI volume space centers on its MNI centroid coordinate and then map the ROI to the Conte69 human surface-based atlas. The ROI on the surface consists of the vertexes in which the voxels belong to the ROI in the volume space. We mapped each sample to the region in which most vertexes belong to in the corresponding ROI. We only analyzed samples in the left brain because among the six brains used to obtain the samples, only 2 had samples in the right brain. The gene expression profiles of each MMP region are the average gene expression of all samples mapped to the region. Brain regions having fewer than 5 AHBA samples mapped to it were filtered out in our subsequent analysis to ensure more reliable results. Finally, 74 regions in the left brain remained after these exclusion criteria were applied.

### Developmental brain gene expression data

To characterize how brain expression of the gene (that are correlated with temporal variability of brain networks) changes with age, we used the developmental brain gene expression profiles in BrainSpan Atlas, also provided by the Allen Brain Atlas. We obtained the processed expression profiles by using the R package ‘ABAData’. In total, 16 brain regions were sampled and analyzed in at least 20 age categories. Expression of a gene for a given brain region and developmental stage are the mean the expression value of all samples.

### Association analysis between temporal variability and gene expression across brain regions

Correlation analysis was performed between temporal variability of brain regions and corresponding gene expression to identify genes/biological pathways correlated to dynamic FC changes. That is, for each of the 20738 genes in the AHBA dataset, correlation analysis between regional gene expression and regional temporal variability was performed across all 74 brain regions. A gene is supposed to be significantly related to temporal variability if the corresponding correlation coefficient can pass the Bonferroni correction (p<0.05/20738) for each of the four fMRI runs. After identifying genes significantly related to temporal variability, we next ^68^ carried out gene enrichment analyses to identify the relevant biological pathways ^68^.

## Acknowledgment

Jie Zhang was supported by the National Natural Science Foundation of China (NSFC 61573107), Special Funds for Major State Basic Research Projects of China (2015CB856003) and National Basic Research Program of China (Precision Psychiatry Program, No. 2016YFC0906402). Jianfeng Feng work was supported by Shanghai Science & Technology Innovation Plan key project Grant Nos. 15JC1400101 and 16JC1420402; National Natural Science Foundation of China Grant Nos. 71661167002, 91630314; Shanghai AI Platform for Diagnosis and Treatment of Brain Diseases Grant No. 2016-17; Base for Introducing Talents of Discipline to Universities Grant No. B18015;

## Supplemental Materials

Supplemental Table 1 The temporal variability for 360 brain regions from HCP_MMP1.0 parcellation. Resting-fMRI data from HCP were used, and group average results were presented.

Supplemental Table 2 The temporal variability of brain regions using AAL2 template from HCP data and UK Biobank data, respectively. A high correlation between regional variability from HCP and UK Biobank was found.

Supplemental Table 3 Genes whose expressions were significantly correlated with dynamic functional connectivity changes in the brain (both positively- and negatively-correlated), Bonfferoni corrected.

Supplemental Table 4 Biological pathways identified for the genes significantly negatively correlated with temporal variability of functional brain networks, Bonfferoni corrected.

Supplemental Table 5 Biological pathways identified for the genes significantly positively correlated with temporal variability of functional brain networks, Bonfferoni corrected.

Supplemental Table 6 Correlation between temporal variability of brain regions and age using Lifespan data from NKIRockland Sample (http://fcon_1000.projects.nitrc.org/indi/pro/nki.html) from the Nathan Kline Institute.

Supplemental Table 7 The 180 areas of the cortical parcellation with index number, short name, description from HCP.

