## Supplementary Materials for "Dynamic brain network changes in resting-state reflect neuroplasticity: molecular and cognitive evidence" (from another version of this preprint: version 1, 2019-07-08)

**Supplementary Table 1**

|  | **Brain region** | **Temporal Variability** |  | **Brain region** | **Temporal Variability** |  | **Brain region** | **Temporal Variability** |
| --- | --- | --- | --- | --- | --- | --- | --- | --- |
| **1** | R_V1_ROI | 0.704206927 | **121** | R_ProS_ROI | 0.819207 | **241** | L_a24_ROI | 0.88305 |
| **2** | R_MST_ROI | 0.719346303 | **122** | R_PeEc_ROI | 0.926591 | **242** | L_d32_ROI | 0.824179 |
| **3** | R_V6_ROI | 0.665503431 | **123** | R_STGa_ROI | 0.868513 | **243** | L_8BM_ROI | 0.794676 |
| **4** | R_V2_ROI | 0.639629811 | **124** | R_PBelt_ROI | 0.797522 | **244** | L_p32_ROI | 0.869007 |
| **5** | R_V3_ROI | 0.667385711 | **125** | R_A5_ROI | 0.77813 | **245** | L_10r_ROI | 0.828862 |
| **6** | R_V4_ROI | 0.658085652 | **126** | R_PHA1_ROI | 0.900826 | **246** | L_47m_ROI | 0.892591 |
| **7** | R_V8_ROI | 0.721812041 | **127** | R_PHA3_ROI | 0.884791 | **247** | L_8Av_ROI | 0.727356 |
| **8** | R_4_ROI | 0.710981645 | **128** | R_STSda_ROI | 0.811022 | **248** | L_8Ad_ROI | 0.774396 |
| **9** | R_3b_ROI | 0.725282288 | **129** | R_STSdp_ROI | 0.804089 | **249** | L_9m_ROI | 0.752874 |
| **10** | R_FEF_ROI | 0.72354115 | **130** | R_STSvp_ROI | 0.771979 | **250** | L_8BL_ROI | 0.763065 |
| **11** | R_PEF_ROI | 0.755777043 | **131** | R_TGd_ROI | 0.80879 | **251** | L_9p_ROI | 0.769934 |
| **12** | R_55b_ROI | 0.823603418 | **132** | R_TE1a_ROI | 0.757168 | **252** | L_10d_ROI | 0.775636 |
| **13** | R_V3A_ROI | 0.633710605 | **133** | R_TE1p_ROI | 0.76141 | **253** | L_8C_ROI | 0.759169 |
| **14** | R_RSC_ROI | 0.860126074 | **134** | R_TE2a_ROI | 0.830622 | **254** | L_44_ROI | 0.822576 |
| **15** | R_POS2_ROI | 0.764199788 | **135** | R_TF_ROI | 0.888881 | **255** | L_45_ROI | 0.795587 |
| **16** | R_V7_ROI | 0.668180508 | **136** | R_TE2p_ROI | 0.79727 | **256** | L_47l_ROI | 0.80049 |
| **17** | R_IPS1_ROI | 0.69324476 | **137** | R_PHT_ROI | 0.725844 | **257** | L_a47r_ROI | 0.792999 |
| **18** | R_FFC_ROI | 0.737440066 | **138** | R_PH_ROI | 0.708722 | **258** | L_6r_ROI | 0.744559 |
| **19** | R_V3B_ROI | 0.662099851 | **139** | R_TPOJ1_ROI | 0.764824 | **259** | L_IFJa_ROI | 0.819014 |
| **20** | R_LO1_ROI | 0.688795759 | **140** | R_TPOJ2_ROI | 0.727632 | **260** | L_IFJp_ROI | 0.813843 |
| **21** | R_LO2_ROI | 0.726901489 | **141** | R_TPOJ3_ROI | 0.75761 | **261** | L_IFSp_ROI | 0.822491 |
| **22** | R_PIT_ROI | 0.740707089 | **142** | R_DVT_ROI | 0.721574 | **262** | L_IFSa_ROI | 0.790921 |
| **23** | R_MT_ROI | 0.749338897 | **143** | R_PGp_ROI | 0.713224 | **263** | L_p9-46v_ROI | 0.770011 |
| **24** | R_A1_ROI | 0.837773449 | **144** | R_IP2_ROI | 0.734678 | **264** | L_46_ROI | 0.71953 |
| **25** | R_PSL_ROI | 0.74191274 | **145** | R_IP1_ROI | 0.739908 | **265** | L_a9-46v_ROI | 0.791465 |
| **26** | R_SFL_ROI | 0.853517488 | **146** | R_IP0_ROI | 0.718815 | **266** | L_9-46d_ROI | 0.784031 |
| **27** | R_PCV_ROI | 0.792327738 | **147** | R_PFop_ROI | 0.714867 | **267** | L_9a_ROI | 0.786364 |
| **28** | R_STV_ROI | 0.775059457 | **148** | R_PF_ROI | 0.672365 | **268** | L_10v_ROI | 0.821206 |
| **29** | R_7Pm_ROI | 0.788532125 | **149** | R_PFm_ROI | 0.714744 | **269** | L_a10p_ROI | 0.831084 |
| **30** | R_7m_ROI | 0.762014761 | **150** | R_PGi_ROI | 0.70638 | **270** | L_10pp_ROI | 0.881435 |
| **31** | R_POS1_ROI | 0.80183118 | **151** | R_PGs_ROI | 0.704817 | **271** | L_11l_ROI | 0.848082 |
| **32** | R_23d_ROI | 0.857561716 | **152** | R_V6A_ROI | 0.696348 | **272** | L_13l_ROI | 0.918233 |
| **33** | R_v23ab_ROI | 0.811662155 | **153** | R_VMV1_ROI | 0.768909 | **273** | L_OFC_ROI | 0.934796 |
| **34** | R_d23ab_ROI | 0.863572234 | **154** | R_VMV3_ROI | 0.794032 | **274** | L_47s_ROI | 0.888423 |
| **35** | R_31pv_ROI | 0.775550105 | **155** | R_PHA2_ROI | 0.907433 | **275** | L_LIPd_ROI | 0.758828 |
| **36** | R_5m_ROI | 0.778228021 | **156** | R_V4t_ROI | 0.715651 | **276** | L_6a_ROI | 0.739136 |
| **37** | R_5mv_ROI | 0.736616367 | **157** | R_FST_ROI | 0.713556 | **277** | L_i6-8_ROI | 0.794245 |
| **38** | R_23c_ROI | 0.7401215 | **158** | R_V3CD_ROI | 0.670743 | **278** | L_s6-8_ROI | 0.837789 |
| **39** | R_5L_ROI | 0.778814321 | **159** | R_LO3_ROI | 0.679435 | **279** | L_43_ROI | 0.776051 |
| **40** | R_24dd_ROI | 0.783509739 | **160** | R_VMV2_ROI | 0.793692 | **280** | L_OP4_ROI | 0.759208 |
| **41** | R_24dv_ROI | 0.814370958 | **161** | R_31pd_ROI | 0.805987 | **281** | L_OP1_ROI | 0.782219 |
| **42** | R_7AL_ROI | 0.750893319 | **162** | R_31a_ROI | 0.818548 | **282** | L_OP2-3_ROI | 0.819317 |
| **43** | R_SCEF_ROI | 0.739165841 | **163** | R_VVC_ROI | 0.790548 | **283** | L_52_ROI | 0.858169 |
| **44** | R_6ma_ROI | 0.761321675 | **164** | R_25_ROI | 0.955196 | **284** | L_RI_ROI | 0.792675 |
| **45** | R_7Am_ROI | 0.707158048 | **165** | R_s32_ROI | 0.893465 | **285** | L_PFcm_ROI | 0.753481 |
| **46** | R_7PL_ROI | 0.734435928 | **166** | R_pOFC_ROI | 0.951019 | **286** | L_PoI2_ROI | 0.85317 |
| **47** | R_7PC_ROI | 0.730858798 | **167** | R_PoI1_ROI | 0.821636 | **287** | L_TA2_ROI | 0.851989 |
| **48** | R_LIPv_ROI | 0.673290115 | **168** | R_Ig_ROI | 0.830303 | **288** | L_FOP4_ROI | 0.778343 |
| **49** | R_VIP_ROI | 0.718176582 | **169** | R_FOP5_ROI | 0.809113 | **289** | L_MI_ROI | 0.849379 |
| **50** | R_MIP_ROI | 0.713855062 | **170** | R_p10p_ROI | 0.792426 | **290** | L_Pir_ROI | 0.942447 |
| **51** | R_1_ROI | 0.7053639 | **171** | R_p47r_ROI | 0.786083 | **291** | L_AVI_ROI | 0.89809 |
| **52** | R_2_ROI | 0.70448089 | **172** | R_TGv_ROI | 0.915889 | **292** | L_AAIC_ROI | 0.952104 |
| **53** | R_3a_ROI | 0.730775908 | **173** | R_MBelt_ROI | 0.86749 | **293** | L_FOP1_ROI | 0.826414 |
| **54** | R_6d_ROI | 0.805747546 | **174** | R_LBelt_ROI | 0.816651 | **294** | L_FOP3_ROI | 0.871675 |
| **55** | R_6mp_ROI | 0.784400158 | **175** | R_A4_ROI | 0.767155 | **295** | L_FOP2_ROI | 0.855116 |
| **56** | R_6v_ROI | 0.757123972 | **176** | R_STSva_ROI | 0.796422 | **296** | L_PFt_ROI | 0.734873 |
| **57** | R_p24pr_ROI | 0.866377071 | **177** | R_TE1m_ROI | 0.783169 | **297** | L_AIP_ROI | 0.716467 |
| **58** | R_33pr_ROI | 0.93681163 | **178** | R_PI_ROI | 0.925185 | **298** | L_EC_ROI | 0.946744 |
| **59** | R_a24pr_ROI | 0.844101154 | **179** | R_a32pr_ROI | 0.826597 | **299** | L_PreS_ROI | 0.910465 |
| **60** | R_p32pr_ROI | 0.772978068 | **180** | R_p24_ROI | 0.894274 | **300** | L_H_ROI | 0.913926 |
| **61** | R_a24_ROI | 0.892852477 | **181** | L_V1_ROI | 0.709077 | **301** | L_ProS_ROI | 0.809805 |
| **62** | R_d32_ROI | 0.826683301 | **182** | L_MST_ROI | 0.726121 | **302** | L_PeEc_ROI | 0.924461 |
| **63** | R_8BM_ROI | 0.791154949 | **183** | L_V6_ROI | 0.669202 | **303** | L_STGa_ROI | 0.882536 |
| **64** | R_p32_ROI | 0.862314005 | **184** | L_V2_ROI | 0.638756 | **304** | L_PBelt_ROI | 0.781486 |
| **65** | R_10r_ROI | 0.83147274 | **185** | L_V3_ROI | 0.650314 | **305** | L_A5_ROI | 0.800559 |
| **66** | R_47m_ROI | 0.872691121 | **186** | L_V4_ROI | 0.665066 | **306** | L_PHA1_ROI | 0.894828 |
| **67** | R_8Av_ROI | 0.740987664 | **187** | L_V8_ROI | 0.734794 | **307** | L_PHA3_ROI | 0.860978 |
| **68** | R_8Ad_ROI | 0.767047411 | **188** | L_4_ROI | 0.706311 | **308** | L_STSda_ROI | 0.831691 |
| **69** | R_9m_ROI | 0.761912115 | **189** | L_3b_ROI | 0.713296 | **309** | L_STSdp_ROI | 0.788747 |
| **70** | R_8BL_ROI | 0.752599384 | **190** | L_FEF_ROI | 0.728445 | **310** | L_STSvp_ROI | 0.758988 |
| **71** | R_9p_ROI | 0.802696251 | **191** | L_PEF_ROI | 0.775856 | **311** | L_TGd_ROI | 0.827121 |
| **72** | R_10d_ROI | 0.786773211 | **192** | L_55b_ROI | 0.815619 | **312** | L_TE1a_ROI | 0.745757 |
| **73** | R_8C_ROI | 0.765774401 | **193** | L_V3A_ROI | 0.632778 | **313** | L_TE1p_ROI | 0.768565 |
| **74** | R_44_ROI | 0.808257032 | **194** | L_RSC_ROI | 0.846529 | **314** | L_TE2a_ROI | 0.860856 |
| **75** | R_45_ROI | 0.820996597 | **195** | L_POS2_ROI | 0.774427 | **315** | L_TF_ROI | 0.913698 |
| **76** | R_47l_ROI | 0.781931875 | **196** | L_V7_ROI | 0.681949 | **316** | L_TE2p_ROI | 0.818243 |
| **77** | R_a47r_ROI | 0.780057289 | **197** | L_IPS1_ROI | 0.688803 | **317** | L_PHT_ROI | 0.748556 |
| **78** | R_6r_ROI | 0.716906383 | **198** | L_FFC_ROI | 0.747446 | **318** | L_PH_ROI | 0.719898 |
| **79** | R_IFJa_ROI | 0.816725185 | **199** | L_V3B_ROI | 0.669104 | **319** | L_TPOJ1_ROI | 0.78622 |
| **80** | R_IFJp_ROI | 0.819449571 | **200** | L_LO1_ROI | 0.701756 | **320** | L_TPOJ2_ROI | 0.748197 |
| **81** | R_IFSp_ROI | 0.814796267 | **201** | L_LO2_ROI | 0.714679 | **321** | L_TPOJ3_ROI | 0.774247 |
| **82** | R_IFSa_ROI | 0.754688786 | **202** | L_PIT_ROI | 0.754052 | **322** | L_DVT_ROI | 0.727101 |
| **83** | R_p9-46v_ROI | 0.751339722 | **203** | L_MT_ROI | 0.743887 | **323** | L_PGp_ROI | 0.725869 |
| **84** | R_46_ROI | 0.717628791 | **204** | L_A1_ROI | 0.834793 | **324** | L_IP2_ROI | 0.736224 |
| **85** | R_a9-46v_ROI | 0.766642479 | **205** | L_PSL_ROI | 0.808359 | **325** | L_IP1_ROI | 0.755685 |
| **86** | R_9-46d_ROI | 0.770368277 | **206** | L_SFL_ROI | 0.834467 | **326** | L_IP0_ROI | 0.716206 |
| **87** | R_9a_ROI | 0.79284859 | **207** | L_PCV_ROI | 0.794564 | **327** | L_PFop_ROI | 0.728728 |
| **88** | R_10v_ROI | 0.797702502 | **208** | L_STV_ROI | 0.787395 | **328** | L_PF_ROI | 0.702656 |
| **89** | R_a10p_ROI | 0.831269876 | **209** | L_7Pm_ROI | 0.78513 | **329** | L_PFm_ROI | 0.708868 |
| **90** | R_10pp_ROI | 0.886952665 | **210** | L_7m_ROI | 0.747829 | **330** | L_PGi_ROI | 0.705066 |
| **91** | R_11l_ROI | 0.834684162 | **211** | L_POS1_ROI | 0.787259 | **331** | L_PGs_ROI | 0.695676 |
| **92** | R_13l_ROI | 0.912048162 | **212** | L_23d_ROI | 0.855787 | **332** | L_V6A_ROI | 0.68615 |
| **93** | R_OFC_ROI | 0.955859452 | **213** | L_v23ab_ROI | 0.79579 | **333** | L_VMV1_ROI | 0.77867 |
| **94** | R_47s_ROI | 0.877225646 | **214** | L_d23ab_ROI | 0.801602 | **334** | L_VMV3_ROI | 0.774329 |
| **95** | R_LIPd_ROI | 0.76492529 | **215** | L_31pv_ROI | 0.76289 | **335** | L_PHA2_ROI | 0.89308 |
| **96** | R_6a_ROI | 0.736112692 | **216** | L_5m_ROI | 0.793222 | **336** | L_V4t_ROI | 0.721926 |
| **97** | R_i6-8_ROI | 0.795302452 | **217** | L_5mv_ROI | 0.752502 | **337** | L_FST_ROI | 0.712296 |
| **98** | R_s6-8_ROI | 0.823893406 | **218** | L_23c_ROI | 0.761943 | **338** | L_V3CD_ROI | 0.682527 |
| **99** | R_43_ROI | 0.754962524 | **219** | L_5L_ROI | 0.784286 | **339** | L_LO3_ROI | 0.691659 |
| **100** | R_OP4_ROI | 0.745866038 | **220** | L_24dd_ROI | 0.790325 | **340** | L_VMV2_ROI | 0.816186 |
| **101** | R_OP1_ROI | 0.802658201 | **221** | L_24dv_ROI | 0.813277 | **341** | L_31pd_ROI | 0.776512 |
| **102** | R_OP2-3_ROI | 0.828872304 | **222** | L_7AL_ROI | 0.730199 | **342** | L_31a_ROI | 0.816686 |
| **103** | R_52_ROI | 0.889460824 | **223** | L_SCEF_ROI | 0.76217 | **343** | L_VVC_ROI | 0.777493 |
| **104** | R_RI_ROI | 0.766548337 | **224** | L_6ma_ROI | 0.781955 | **344** | L_25_ROI | 0.939208 |
| **105** | R_PFcm_ROI | 0.739176359 | **225** | L_7Am_ROI | 0.714801 | **345** | L_s32_ROI | 0.898948 |
| **106** | R_PoI2_ROI | 0.825108001 | **226** | L_7PL_ROI | 0.731132 | **346** | L_pOFC_ROI | 0.956395 |
| **107** | R_TA2_ROI | 0.870573698 | **227** | L_7PC_ROI | 0.728011 | **347** | L_PoI1_ROI | 0.852809 |
| **108** | R_FOP4_ROI | 0.764852205 | **228** | L_LIPv_ROI | 0.685703 | **348** | L_Ig_ROI | 0.885422 |
| **109** | R_MI_ROI | 0.812499712 | **229** | L_VIP_ROI | 0.720408 | **349** | L_FOP5_ROI | 0.841256 |
| **110** | R_Pir_ROI | 0.926450196 | **230** | L_MIP_ROI | 0.713926 | **350** | L_p10p_ROI | 0.833425 |
| **111** | R_AVI_ROI | 0.840359895 | **231** | L_1_ROI | 0.700844 | **351** | L_p47r_ROI | 0.796967 |
| **112** | R_AAIC_ROI | 0.943901673 | **232** | L_2_ROI | 0.701479 | **352** | L_TGv_ROI | 0.916813 |
| **113** | R_FOP1_ROI | 0.821768936 | **233** | L_3a_ROI | 0.727546 | **353** | L_MBelt_ROI | 0.84762 |
| **114** | R_FOP3_ROI | 0.845574339 | **234** | L_6d_ROI | 0.800412 | **354** | L_LBelt_ROI | 0.822498 |
| **115** | R_FOP2_ROI | 0.842771908 | **235** | L_6mp_ROI | 0.771505 | **355** | L_A4_ROI | 0.760843 |
| **116** | R_PFt_ROI | 0.740317572 | **236** | L_6v_ROI | 0.778636 | **356** | L_STSva_ROI | 0.776968 |
| **117** | R_AIP_ROI | 0.7069345 | **237** | L_p24pr_ROI | 0.88889 | **357** | L_TE1m_ROI | 0.78994 |
| **118** | R_EC_ROI | 0.954606337 | **238** | L_33pr_ROI | 0.928784 | **358** | L_PI_ROI | 0.88656 |
| **119** | R_PreS_ROI | 0.922325338 | **239** | L_a24pr_ROI | 0.813188 | **359** | L_a32pr_ROI | 0.850997 |
| **120** | R_H_ROI | 0.923378974 | **240** | L_p32pr_ROI | 0.780401 | **360** | L_p24_ROI | 0.881842 |

**Supplemental Table 2** Genes whose expressions were significantly correlated with dynamic functional connectivity changes in the brain (both positively- and negatively-correlated), Bonfferoni corrected.

| **The temporal variability of HCP and Biobank (AAL2)** | | | | | | | |  |
| --- | --- | --- | --- | --- | --- | --- | --- | --- |
| **Region Names** | **HCP_Run_1** | **HCP_Run_2** | **HCP_Run_3** | **HCP_Run_4** | **HCP_mean** | **HCP_rank** | **Biobank** | **Biobank_rank** |
| Olfactory_R | 0.9260 | 0.9220 | 0.9277 | 0.9240 | 0.9249 | 2 | 0.9059 | 1 |
| Pallidum_L | 0.9233 | 0.9307 | 0.9302 | 0.9278 | 0.9280 | 1 | 0.8994 | 2 |
| Olfactory_L | 0.9006 | 0.9217 | 0.9072 | 0.9170 | 0.9116 | 6 | 0.8988 | 3 |
| Amygdala_R | 0.9086 | 0.9149 | 0.9191 | 0.9055 | 0.9120 | 5 | 0.8979 | 4 |
| Amygdala_L | 0.9006 | 0.9193 | 0.9147 | 0.9104 | 0.9113 | 8 | 0.8963 | 5 |
| Pallidum_R | 0.9163 | 0.9154 | 0.9246 | 0.9141 | 0.9176 | 3 | 0.8863 | 6 |
| Hippocampus_R | 0.8724 | 0.8692 | 0.8845 | 0.8692 | 0.8738 | 16 | 0.8837 | 7 |
| OFCmed_L | 0.9072 | 0.9201 | 0.9136 | 0.9135 | 0.9136 | 4 | 0.8832 | 8 |
| ParaHippocampal_L | 0.8620 | 0.8824 | 0.8727 | 0.8799 | 0.8743 | 14 | 0.8764 | 9 |
| Hippocampus_L | 0.8622 | 0.8727 | 0.8743 | 0.8727 | 0.8705 | 19 | 0.8752 | 10 |
| Caudate_L | 0.8842 | 0.8917 | 0.8954 | 0.8855 | 0.8892 | 10 | 0.8740 | 11 |
| OFCmed_R | 0.9099 | 0.9126 | 0.9156 | 0.9078 | 0.9115 | 7 | 0.8732 | 12 |
| Caudate_R | 0.8751 | 0.8920 | 0.8867 | 0.8859 | 0.8849 | 11 | 0.8647 | 13 |
| OFCpost_L | 0.8493 | 0.8516 | 0.8631 | 0.8521 | 0.8541 | 24 | 0.8616 | 14 |
| ParaHippocampal_R | 0.8608 | 0.8575 | 0.8741 | 0.8472 | 0.8599 | 21 | 0.8606 | 15 |
| OFCant_L | 0.8824 | 0.9021 | 0.8901 | 0.8951 | 0.8924 | 9 | 0.8603 | 16 |
| OFCpost_R | 0.8426 | 0.8557 | 0.8558 | 0.8536 | 0.8519 | 26 | 0.8578 | 17 |
| Temporal_Pole_Mid_L | 0.8494 | 0.8342 | 0.8579 | 0.8371 | 0.8447 | 28 | 0.8574 | 18 |
| OFClat_R | 0.8497 | 0.8533 | 0.8535 | 0.8576 | 0.8536 | 25 | 0.8572 | 19 |
| Temporal_Pole_Sup_L | 0.8171 | 0.8356 | 0.8405 | 0.8187 | 0.8280 | 38 | 0.8535 | 20 |
| Rectus_R | 0.8790 | 0.8814 | 0.8828 | 0.8835 | 0.8817 | 12 | 0.8523 | 21 |
| Temporal_Pole_Sup_R | 0.8069 | 0.8356 | 0.8341 | 0.8121 | 0.8222 | 41 | 0.8498 | 22 |
| Heschl_L | 0.7923 | 0.8084 | 0.8206 | 0.7864 | 0.8019 | 61 | 0.8491 | 23 |
| Temporal_Pole_Mid_R | 0.8230 | 0.8342 | 0.8341 | 0.8331 | 0.8311 | 35 | 0.8490 | 24 |
| OFClat_L | 0.8427 | 0.8440 | 0.8475 | 0.8487 | 0.8457 | 27 | 0.8457 | 25 |
| OFCant_R | 0.8716 | 0.8742 | 0.8762 | 0.8741 | 0.8740 | 15 | 0.8447 | 26 |
| Cingulate_Post_R | 0.8594 | 0.8481 | 0.8667 | 0.8540 | 0.8571 | 23 | 0.8393 | 27 |
| Putamen_L | 0.8747 | 0.8801 | 0.8802 | 0.8777 | 0.8782 | 13 | 0.8374 | 28 |
| Temporal_Inf_L | 0.8337 | 0.8423 | 0.8506 | 0.8294 | 0.8390 | 31 | 0.8368 | 29 |
| Thalamus_L | 0.8691 | 0.8689 | 0.8801 | 0.8669 | 0.8713 | 18 | 0.8362 | 30 |
| Thalamus_R | 0.8636 | 0.8633 | 0.8739 | 0.8582 | 0.8648 | 20 | 0.8323 | 31 |
| Frontal_Inf_Orb_2_R | 0.8253 | 0.8430 | 0.8396 | 0.8396 | 0.8369 | 33 | 0.8297 | 32 |
| Temporal_Inf_R | 0.8105 | 0.8374 | 0.8336 | 0.8245 | 0.8265 | 39 | 0.8280 | 33 |
| Putamen_R | 0.8717 | 0.8697 | 0.8785 | 0.8684 | 0.8721 | 17 | 0.8277 | 34 |
| Frontal_Inf_Orb_2_L | 0.8287 | 0.8264 | 0.8420 | 0.8227 | 0.8300 | 37 | 0.8272 | 35 |
| Heschl_R | 0.7839 | 0.8093 | 0.8147 | 0.7837 | 0.7979 | 65 | 0.8232 | 36 |
| Cingulate_Post_L | 0.8316 | 0.8315 | 0.8386 | 0.8420 | 0.8359 | 34 | 0.8209 | 37 |
| Rectus_L | 0.8482 | 0.8632 | 0.8573 | 0.8661 | 0.8587 | 22 | 0.8194 | 38 |
| Paracentral_Lobule_R | 0.7930 | 0.8174 | 0.8189 | 0.8057 | 0.8087 | 55 | 0.8180 | 39 |
| Frontal_Med_Orb_R | 0.8166 | 0.8126 | 0.8200 | 0.8325 | 0.8204 | 42 | 0.8174 | 40 |
| Cingulate_Ant_R | 0.8364 | 0.8436 | 0.8462 | 0.8438 | 0.8425 | 30 | 0.8130 | 41 |
| Cingulate_Ant_L | 0.8360 | 0.8452 | 0.8504 | 0.8426 | 0.8435 | 29 | 0.8102 | 42 |
| Frontal_Inf_Tri_L | 0.8195 | 0.8264 | 0.8333 | 0.8200 | 0.8248 | 40 | 0.8094 | 43 |
| Paracentral_Lobule_L | 0.7942 | 0.8210 | 0.8205 | 0.8071 | 0.8107 | 53 | 0.8092 | 44 |
| Frontal_Inf_Tri_R | 0.8207 | 0.8322 | 0.8374 | 0.8321 | 0.8306 | 36 | 0.8054 | 45 |
| Temporal_Mid_L | 0.7947 | 0.8052 | 0.8166 | 0.7965 | 0.8033 | 60 | 0.8045 | 46 |
| Frontal_Med_Orb_L | 0.8096 | 0.7963 | 0.8123 | 0.8189 | 0.8093 | 54 | 0.8040 | 47 |
| Occipital_Inf_R | 0.8261 | 0.8454 | 0.8442 | 0.8390 | 0.8387 | 32 | 0.8000 | 48 |
| Temporal_Sup_R | 0.7691 | 0.8066 | 0.8049 | 0.7726 | 0.7883 | 69 | 0.7989 | 49 |
| Temporal_Mid_R | 0.7997 | 0.8213 | 0.8214 | 0.8071 | 0.8124 | 52 | 0.7980 | 50 |
| Frontal_Inf_Oper_L | 0.8049 | 0.8022 | 0.8147 | 0.8066 | 0.8071 | 58 | 0.7963 | 51 |
| Precuneus_L | 0.8069 | 0.8147 | 0.8198 | 0.8116 | 0.8132 | 50 | 0.7955 | 52 |
| Parietal_Inf_R | 0.8196 | 0.8166 | 0.8237 | 0.8207 | 0.8202 | 43 | 0.7945 | 53 |
| Angular_R | 0.8155 | 0.8142 | 0.8226 | 0.8204 | 0.8182 | 45 | 0.7941 | 54 |
| Frontal_Mid_2_L | 0.8118 | 0.8204 | 0.8212 | 0.8183 | 0.8179 | 46 | 0.7941 | 55 |
| Frontal_Sup_2_L | 0.8119 | 0.8182 | 0.8260 | 0.8182 | 0.8186 | 44 | 0.7918 | 56 |
| Cingulate_Mid_L | 0.8052 | 0.8089 | 0.8192 | 0.7962 | 0.8074 | 57 | 0.7908 | 57 |
| Angular_L | 0.8012 | 0.7875 | 0.8068 | 0.8080 | 0.8009 | 62 | 0.7907 | 58 |
| Frontal_Sup_2_R | 0.8099 | 0.8164 | 0.8218 | 0.8171 | 0.8163 | 49 | 0.7894 | 59 |
| Precuneus_R | 0.8142 | 0.8149 | 0.8258 | 0.8109 | 0.8165 | 48 | 0.7873 | 60 |
| Parietal_Inf_L | 0.8079 | 0.8158 | 0.8151 | 0.8142 | 0.8132 | 51 | 0.7873 | 61 |
| Frontal_Sup_Medial_R | 0.7947 | 0.7913 | 0.8036 | 0.8053 | 0.7987 | 64 | 0.7872 | 62 |
| Cingulate_Mid_R | 0.8025 | 0.8076 | 0.8150 | 0.7970 | 0.8055 | 59 | 0.7837 | 63 |
| Frontal_Mid_2_R | 0.8045 | 0.8053 | 0.8122 | 0.8083 | 0.8076 | 56 | 0.7825 | 64 |
| Supp_Motor_Area_L | 0.7851 | 0.7980 | 0.8057 | 0.7941 | 0.7957 | 67 | 0.7790 | 65 |
| Occipital_Inf_L | 0.8066 | 0.8204 | 0.8267 | 0.8152 | 0.8172 | 47 | 0.7783 | 66 |
| Frontal_Sup_Medial_L | 0.7903 | 0.7884 | 0.8020 | 0.8006 | 0.7953 | 68 | 0.7783 | 67 |
| Temporal_Sup_L | 0.7475 | 0.7740 | 0.7785 | 0.7479 | 0.7620 | 84 | 0.7754 | 68 |
| Fusiform_R | 0.7735 | 0.7951 | 0.7972 | 0.7836 | 0.7874 | 70 | 0.7728 | 69 |
| Frontal_Inf_Oper_R | 0.7873 | 0.7901 | 0.8011 | 0.8058 | 0.7961 | 66 | 0.7690 | 70 |
| Fusiform_L | 0.7843 | 0.8085 | 0.8088 | 0.7969 | 0.7996 | 63 | 0.7682 | 71 |
| Insula_L | 0.7636 | 0.7696 | 0.7805 | 0.7706 | 0.7711 | 79 | 0.7647 | 72 |
| Precentral_L | 0.7550 | 0.7713 | 0.7776 | 0.7700 | 0.7685 | 81 | 0.7625 | 73 |
| SupraMarginal_L | 0.7750 | 0.7728 | 0.7879 | 0.7746 | 0.7776 | 77 | 0.7619 | 74 |
| Supp_Motor_Area_R | 0.7625 | 0.7768 | 0.7822 | 0.7758 | 0.7743 | 78 | 0.7614 | 75 |
| Parietal_Sup_L | 0.7783 | 0.7868 | 0.7889 | 0.7888 | 0.7857 | 71 | 0.7610 | 76 |
| Rolandic_Oper_L | 0.7296 | 0.7382 | 0.7547 | 0.7315 | 0.7385 | 93 | 0.7585 | 77 |
| Parietal_Sup_R | 0.7749 | 0.7822 | 0.7868 | 0.7870 | 0.7827 | 72 | 0.7574 | 78 |
| Insula_R | 0.7563 | 0.7583 | 0.7752 | 0.7594 | 0.7623 | 83 | 0.7544 | 79 |
| Postcentral_R | 0.7388 | 0.7662 | 0.7682 | 0.7523 | 0.7564 | 85 | 0.7543 | 80 |
| Calcarine_L | 0.7731 | 0.7862 | 0.7846 | 0.7759 | 0.7800 | 75 | 0.7540 | 81 |
| Cuneus_L | 0.7518 | 0.7701 | 0.7698 | 0.7577 | 0.7624 | 82 | 0.7535 | 82 |
| Postcentral_L | 0.7340 | 0.7631 | 0.7639 | 0.7453 | 0.7516 | 88 | 0.7528 | 83 |
| Calcarine_R | 0.7714 | 0.7818 | 0.7832 | 0.7761 | 0.7781 | 76 | 0.7527 | 84 |
| Precentral_R | 0.7340 | 0.7563 | 0.7593 | 0.7514 | 0.7503 | 90 | 0.7523 | 85 |
| SupraMarginal_R | 0.7659 | 0.7615 | 0.7758 | 0.7736 | 0.7692 | 80 | 0.7518 | 86 |
| Rolandic_Oper_R | 0.7212 | 0.7417 | 0.7465 | 0.7346 | 0.7360 | 94 | 0.7474 | 87 |
| Occipital_Mid_R | 0.7701 | 0.7852 | 0.7851 | 0.7818 | 0.7805 | 74 | 0.7461 | 88 |
| Lingual_R | 0.7427 | 0.7571 | 0.7583 | 0.7496 | 0.7519 | 87 | 0.7401 | 89 |
| Lingual_L | 0.7421 | 0.7584 | 0.7589 | 0.7510 | 0.7526 | 86 | 0.7395 | 90 |
| Occipital_Sup_R | 0.7406 | 0.7515 | 0.7561 | 0.7529 | 0.7503 | 89 | 0.7380 | 91 |
| Cuneus_R | 0.7334 | 0.7479 | 0.7480 | 0.7427 | 0.7430 | 91 | 0.7361 | 92 |
| Occipital_Mid_L | 0.7698 | 0.7864 | 0.7871 | 0.7796 | 0.7808 | 73 | 0.7352 | 93 |
| Occipital_Sup_L | 0.7327 | 0.7422 | 0.7469 | 0.7435 | 0.7413 | 92 | 0.7305 | 94 |
| **Correlation with Biobank results** | | | | | |  |  |  |
|  | 0.9291 | 0.9291 | 0.9479 | 0.9230 | 0.9400 |  |  |  |

**Supplemental Table 3**

| **Negatively related gene** | **R** | **Negatively related gene** | **R** | **Positively related gene** | **R** |
| --- | --- | --- | --- | --- | --- |
| A26C1B | -0.623534687 | LMLN | -0.563927051 | A1BG | 0.65924 |
| ABCC8 | -0.587674615 | LMOD3 | -0.58819791 | ABCA17P | 0.57706 |
| ABTB1 | -0.633960825 | LOC100128174 | -0.568924953 | ACOT12 | 0.63173 |
| AC084125.1 | -0.633385838 | LOC100133150 | -0.658869771 | ADCYAP1R1 | 0.57573 |
| ACSL3 | -0.60950679 | LOC100289408 | -0.55764426 | ADIPOR1 | 0.57585 |
| ACTR1B | -0.672769454 | LOC407835 | -0.549769451 | AK3L1 | 0.60774 |
| AFAP1L1 | -0.591826891 | LOC440459 | -0.632751686 | APOBEC3F | 0.57158 |
| AGPAT9 | -0.63698093 | LOC642513 | -0.666880598 | APOE | 0.56293 |
| ALDH1A3 | -0.701598514 | LRRC38 | -0.595787972 | ARHGAP6 | 0.58001 |
| AMMECR1 | -0.60804411 | LRRC4 | -0.608498058 | C10orf105 | 0.55348 |
| ANKH | -0.657307118 | LUZP1 | -0.605506017 | C20orf26 | 0.59299 |
| ANKRD34C | -0.585229804 | LYSMD4 | -0.659923401 | C2orf73 | 0.57327 |
| ANXA6 | -0.529988319 | MADCAM1 | -0.613051871 | C3orf26 | 0.59174 |
| APOL2 | -0.656761483 | MAFB | -0.571068664 | C4A | 0.55994 |
| AR | -0.598434966 | MAP3K13 | -0.69728066 | C7orf10 | 0.5509 |
| ARHGAP26 | -0.585267915 | MAP9 | -0.682838043 | C8orf30B | 0.57967 |
| ARHGAP9 | -0.605182965 | MAT2B | -0.641495133 | C9orf135 | 0.61672 |
| ARHGEF17 | -0.586753304 | MCF2L2 | -0.540643144 | CADM1 | 0.5603 |
| ARHGEF18 | -0.609989154 | MIR133A1 | -0.576519553 | CBFB | 0.5571 |
| ARL4C | -0.603351374 | MIR31HG | -0.623682467 | CCDC112 | 0.56719 |
| ARMC8 | -0.597202572 | MKX | -0.541819593 | CCDC19 | 0.59752 |
| ARNTL | -0.602707459 | MOB2 | -0.584233974 | CHODL | 0.53712 |
| ASB13 | -0.715955709 | MOGAT1 | -0.62221679 | CHST9 | 0.54516 |
| ATP1A1 | -0.688837255 | MPP1 | -0.681434273 | CKS1B | 0.5379 |
| ATP1A3 | -0.605046837 | MTP18 | -0.576130299 | CNTNAP3 | 0.5756 |
| ATP2B2 | -0.645739983 | MTUS2 | -0.653842594 | COCH | 0.58832 |
| ATP2B3 | -0.597672486 | MYC | -0.618591132 | CPE | 0.55633 |
| AVPI1 | -0.569056509 | MYH7B | -0.64021741 | CPNE2 | 0.56216 |
| BAG5 | -0.548795768 | MYO15A | -0.65629147 | CPNE3 | 0.55357 |
| BCL3 | -0.578556978 | NAT8L | -0.567265192 | CPNE6 | 0.6867 |
| BMP4 | -0.588254902 | NCKIPSD | -0.561915231 | DAK | 0.63934 |
| C10orf140 | -0.744715216 | NEFH | -0.598111993 | DAPL1 | 0.6028 |
| C14orf91 | -0.581375386 | NFIC | -0.679873489 | DCAKD | 0.58443 |
| C17orf75 | -0.587248981 | NIPAL2 | -0.630872569 | DERL1 | 0.56246 |
| C1orf133 | -0.605494213 | NKAPL | -0.621994877 | DHDH | 0.61333 |
| C1orf201 | -0.665256324 | NT5M | -0.699797587 | DNAH12 | 0.56192 |
| C1orf222 | -0.577247065 | OR2AK2 | -0.551875212 | DNAH2 | 0.57033 |
| C1orf71 | -0.562655203 | OR2L8 | -0.613046931 | DNAJC12 | 0.54015 |
| C1RL | -0.606195602 | OSBPL6 | -0.695341484 | DSEL | 0.60278 |
| C20orf39 | -0.581918013 | P2RX6 | -0.588529974 | DTWD1 | 0.54105 |
| C5orf53 | -0.602813044 | PELI3 | -0.663110681 | ECHDC3 | 0.6408 |
| C6orf106 | -0.612036811 | PER1 | -0.608022331 | EDNRB | 0.6087 |
| C6orf136 | -0.545328902 | PFAS | -0.566732059 | EFEMP1 | 0.59433 |
| C6orf211 | -0.63118137 | PGP | -0.552900385 | EIF4EBP1 | 0.59001 |
| C9orf45 | -0.698444177 | PHYH | -0.674586832 | EPCAM | 0.5318 |
| CA13 | -0.588904994 | PLCXD1 | -0.605524661 | ERBB2 | 0.57294 |
| CABP1 | -0.547769595 | PLEKHA6 | -0.661135768 | FABP5P3 | 0.56468 |
| CACNG2 | -0.547755629 | PLEKHM2 | -0.639888732 | FAM181A | 0.57934 |
| CACNG8 | -0.572408355 | PLXDC1 | -0.753451365 | FAM183A | 0.6448 |
| CAMK2G | -0.579301272 | POU6F1 | -0.733030531 | FAM196A | 0.56539 |
| CAMKK1 | -0.536881608 | PPAPDC3 | -0.608433457 | FBXO15 | 0.5502 |
| CAMKK2 | -0.539560659 | PPARGC1A | -0.685489558 | FZD1 | 0.57089 |
| CAMTA2 | -0.611000915 | PPM1A | -0.592982982 | GABRQ | 0.57592 |
| CAPS2 | -0.589280779 | PPP1R28 | -0.53378168 | GNG10 | 0.57214 |
| CCBL1 | -0.60928238 | PPP3CA | -0.57947586 | GOLT1A | 0.56321 |
| CCDC64B | -0.542245871 | PPTC7 | -0.595092416 | GTF2F2 | 0.60253 |
| CCNB1IP1 | -0.623022982 | PRC1 | -0.545155393 | GTF2IRD2P1 | 0.5466 |
| CCNI | -0.668734081 | PRDM1 | -0.565299236 | HCST | 0.62185 |
| CD99L2 | -0.55989084 | PREP | -0.611470847 | HEBP1 | 0.56697 |
| CDS1 | -0.61421565 | PRSS16 | -0.563544902 | HEYL | 0.59282 |
| CHAF1A | -0.59896287 | PVALB | -0.631320742 | HIST1H2BC | 0.62156 |
| CITED2 | -0.569853918 | QRICH2 | -0.570776381 | HIST1H2BH | 0.53684 |
| CLEC16A | -0.621005351 | RAB11FIP3 | -0.556393066 | HIST1H2BO | 0.54122 |
| CLN6 | -0.580886898 | RAB40C | -0.554362806 | HNMT | 0.54737 |
| CNTNAP1 | -0.575594022 | RAD54B | -0.721724711 | IFI27L2 | 0.59133 |
| CORO6 | -0.584077518 | RAI1 | -0.557155902 | IGFBP5 | 0.56621 |
| COX7A1 | -0.566135442 | RCAN2 | -0.597931744 | IL1B | 0.5769 |
| CPEB3 | -0.547223044 | RCN3 | -0.586894174 | KCNJ1 | 0.59989 |
| CPLX1 | -0.640827333 | RELL2 | -0.669030829 | KCTD12 | 0.55955 |
| CPNE9 | -0.567880923 | REPIN1 | -0.610264578 | KLHL13 | 0.59668 |
| CRHR2 | -0.612667389 | RET | -0.626615321 | LECT1 | 0.54971 |
| CTNNAL1 | -0.561615824 | RFPL1 | -0.645799531 | LINC00260 | 0.63552 |
| CTSA | -0.573797776 | RFPL3 | -0.614751196 | LOC100133211 | 0.59039 |
| CUGBP2 | -0.656221576 | RFX5 | -0.693809932 | LOC100292909 | 0.53065 |
| CYP2E1 | -0.53727034 | RHBDL3 | -0.64822942 | LOC653051 | 0.58027 |
| DAPK3 | -0.641888421 | RIMKLA | -0.640865384 | LOC653513 | 0.56414 |
| DCBLD2 | -0.602383849 | RIMS3 | -0.54722649 | LRCH2 | 0.57102 |
| DCLRE1A | -0.673899529 | ROCK2 | -0.641278445 | LRRC36 | 0.63691 |
| DDHD2 | -0.632178502 | ROS1 | -0.533404318 | LRRC3B | 0.60466 |
| DDX3Y | -0.57072256 | RP5-1022P6.1 | -0.619556164 | LRRC56 | 0.63401 |
| DEPDC5 | -0.643937104 | RP5-1068E13.3 | -0.603637267 | MACROD2 | 0.58072 |
| DEXI | -0.637196314 | RPP25 | -0.552097828 | MASP1 | 0.53324 |
| DFNB59 | -0.605110924 | RPS6KA3 | -0.631493574 | MATN2 | 0.58806 |
| DHX35 | -0.681398769 | RRS1 | -0.571409137 | MT1F | 0.61191 |
| DIRAS1 | -0.603769674 | RTKN2 | -0.569786241 | NKAIN4 | 0.66107 |
| DKK1 | -0.627958419 | RUSC2 | -0.532139676 | NUPR1 | 0.5672 |
| DMKN | -0.649351082 | SAT2 | -0.611543932 | OPHN1 | 0.58019 |
| DMRT3 | -0.648164161 | SCN1A | -0.689002844 | PARD3B | 0.57775 |
| DPP4 | -0.614275317 | SCN1B | -0.714484709 | PBEF1 | 0.55868 |
| DPP8 | -0.627173371 | SCN8A | -0.535838588 | PCSK5 | 0.58736 |
| DQX1 | -0.683431222 | SCRT1 | -0.727823793 | PDYN | 0.54723 |
| ECM1 | -0.611213011 | SDHA | -0.60566064 | PEG10 | 0.57637 |
| ECSIT | -0.58662373 | SEMA7A | -0.671377609 | PGAP1 | 0.63222 |
| EFNA5 | -0.621592234 | SERTAD4 | -0.691756307 | PGRMC1 | 0.59025 |
| EHD2 | -0.629605009 | SGCG | -0.573908799 | PKIA | 0.63854 |
| EIF4A2 | -0.588542047 | SHD | -0.747447282 | PLEKHC1 | 0.58693 |
| EIF4E1B | -0.611918485 | SIRT4 | -0.577032093 | PON3 | 0.66339 |
| EIF5A2 | -0.58772877 | SIRT5 | -0.584921523 | PRKG1 | 0.5522 |
| EME1 | -0.63217299 | SIX4 | -0.66776585 | PRMT6 | 0.58901 |
| EMILIN3 | -0.570102902 | SLC12A5 | -0.668507124 | PTPRA | 0.5943 |
| EML6 | -0.570313281 | SLC16A6 | -0.643411686 | RAB8A | 0.54375 |
| ENTPD4 | -0.625634917 | SLC16A7 | -0.687620131 | RALGPS2 | 0.55351 |
| EPN3 | -0.662692408 | SLC25A12 | -0.675254039 | RAVER2 | 0.60153 |
| ESRRA | -0.610573413 | SLC25A5 | -0.59733274 | RNASEH2A | 0.64146 |
| ESYT1 | -0.574328383 | SLC38A11 | -0.569946021 | RP11-45B20.3 | 0.64388 |
| ETV4 | -0.643351545 | SLC39A14 | -0.702743865 | RP11-60I3.5 | 0.5719 |
| EXTL2 | -0.620886017 | SLC45A4 | -0.568124968 | RRAGB | 0.58216 |
| FAM123B | -0.644743684 | SLC5A4 | -0.542669666 | RRAS2 | 0.58024 |
| FAM131B | -0.637334762 | SLC5A6 | -0.628428246 | RSPO3 | 0.56098 |
| FAM83D | -0.66473325 | SLC7A5P2 | -0.574677491 | RYR1 | 0.60953 |
| FBLN7 | -0.666692219 | SLC9A1 | -0.660979522 | RYR3 | 0.58192 |
| FBXO28 | -0.568018585 | SLCO4A1 | -0.73570901 | SAG | 0.6165 |
| FBXO33 | -0.700280121 | SMPX | -0.58461251 | SCIN | 0.61764 |
| FBXO9 | -0.556205948 | SNAP25 | -0.629799201 | SEL1L2 | 0.60749 |
| FCHSD1 | -0.669530226 | SNCB | -0.570477487 | SEMA3D | 0.5694 |
| FES | -0.590536215 | SORL1 | -0.65833077 | SLA | 0.64878 |
| FGF18 | -0.608480099 | SOST | -0.543361996 | SLC2A12 | 0.58408 |
| FGF9 | -0.590585097 | SPAG4 | -0.724581254 | SLC39A6 | 0.59538 |
| FIZ1 | -0.603845165 | SPAG5 | -0.58676178 | SLC4A3 | 0.58818 |
| FNDC4 | -0.581232916 | SPTSSB | -0.622140296 | SLIT3 | 0.62117 |
| FUT11 | -0.55477398 | SRL | -0.613930261 | SMARCD3 | 0.63716 |
| GABRA1 | -0.572413323 | SRPK1 | -0.628976521 | SPAG17 | 0.54657 |
| GABRG2 | -0.525713707 | SSX2IP | -0.532379415 | TDRD5 | 0.60967 |
| GDPD1 | -0.600747674 | ST3GAL3 | -0.56050889 | TMEM176A | 0.59185 |
| GK | -0.655622228 | ST8SIA5 | -0.552161656 | TRAIP | 0.57395 |
| GLB1L3 | -0.617273306 | STAC2 | -0.661490192 | TRIM34 | 0.55918 |
| GLCCI1 | -0.599839236 | STAMBPL1 | -0.525441726 | TSPAN2 | 0.54788 |
| GLS2 | -0.607132887 | STARD5 | -0.631981065 | UCHL3 | 0.64589 |
| GON4L | -0.56335795 | STEAP2 | -0.640256712 | VANGL2 | 0.54642 |
| GPR161 | -0.628968774 | STRBP | -0.631603713 | WFS1 | 0.62084 |
| GRIN2A | -0.57600002 | STS | -0.566543987 | ZMAT1 | 0.53911 |
| GSTT1 | -0.605769709 | STX19 | -0.576982207 | ZNF229 | 0.57226 |
| HAGHL | -0.597129696 | SV2C | -0.599337581 | ZNF311 | 0.606 |
| HAPLN4 | -0.687698636 | SYCP2 | -0.613439966 | ZNF572 | 0.55035 |
| HCN1 | -0.53467872 | SYT12 | -0.575495617 | ZNF706 | 0.56673 |
| HCRP1 | -0.557737796 | TAF4B | -0.579371809 |  |  |
| HECW1 | -0.585214844 | TAPBP | -0.54329996 |  |  |
| HEXIM1 | -0.598816969 | TAS2R9 | -0.541758385 |  |  |
| HIVEP2 | -0.612210513 | TCTEX1D1 | -0.643511737 |  |  |
| HR | -0.725801797 | TDRD1 | -0.594567913 |  |  |
| HSPB6 | -0.567155469 | THAP10 | -0.65165572 |  |  |
| IFFO1 | -0.685611407 | TIAM2 | -0.621228505 |  |  |
| IGFBP2 | -0.539763565 | TMEM170B | -0.592376858 |  |  |
| INA | -0.655298495 | TMEM25 | -0.551387527 |  |  |
| IQSEC3 | -0.581202462 | TMEM81 | -0.690505555 |  |  |
| IRS1 | -0.592722804 | TNIP1 | -0.580770243 |  |  |
| ITPR1 | -0.68568841 | TPTE2P6 | -0.718813418 |  |  |
| JDP2 | -0.611139209 | TRERF1 | -0.61367211 |  |  |
| KCNA2 | -0.703297254 | UCHL5 | -0.598162325 |  |  |
| KCNA5 | -0.654382908 | VAMP1 | -0.59486777 |  |  |
| KCNAB3 | -0.602743885 | YOD1 | -0.608551164 |  |  |
| KCNH1 | -0.566224515 | ZBTB16 | -0.584778898 |  |  |
| KCNJ11 | -0.575483171 | ZBTB7A | -0.554434716 |  |  |
| KCTD9 | -0.585638497 | ZNF25 | -0.609580098 |  |  |
| KIAA0427 | -0.608234753 | ZNF295 | -0.644631605 |  |  |
| KIAA0430 | -0.619736256 | ZNF365 | -0.541342841 |  |  |
| KIAA1107 | -0.574494556 | ZNF641 | -0.601953489 |  |  |
| KIAA1199 | -0.574557841 | ZYX | -0.68006963 |  |  |
| KLC2 | -0.582307058 |  |  |  |  |
| KLF12 | -0.671397496 |  |  |  |  |
| KLHL17 | -0.555136268 |  |  |  |  |
| KRTDAP | -0.580855224 |  |  |  |  |
| LAG3 | -0.613429301 |  |  |  |  |
| LAPTM4B | -0.562245788 |  |  |  |  |
| LEPROTL1 | -0.591728245 |  |  |  |  |
| LINC00515 | -0.654604231 |  |  |  |  |

**Supplemental Table 4**

| **ID** | **Biological Pathway** | | **p-value** | **q-value Bonferroni** | **q-value FDR B&H** | **q-value FDR B&Y** | **Hit in Query List** | | **Hit in Genome** |
| --- | --- | --- | --- | --- | --- | --- | --- | --- | --- |
| **set1** | **synaptic transmission** | | | | | | | | |
| GO:0035637 | multicellular organismal signaling | 1.82E-08 | | 7.30E-05 | 7.30E-05 | 0.000648 | 16 | 197 | |
| GO:0006811 | ion transport | 8.11E-08 | | 0.000325 | 0.000163 | 0.00144 | 50 | 1627 | |
| GO:0030001 | metal ion transport | 3.87E-07 | | 0.00155 | 0.000518 | 0.0046 | 32 | 855 | |
| GO:0006812 | cation transport | 5.41E-06 | | 0.0217 | 0.00376 | 0.0334 | 35 | 1110 | |
| GO:0034220 | ion transmembrane transport | 5.62E-06 | | 0.0226 | 0.00376 | 0.0334 | 34 | 1064 | |
| GO:0019226 | transmission of nerve impulse | 1.58E-05 | | 6.33E-02 | 9.05E-03 | 8.03E-02 | 8 | 80 | |
| GO:0043269 | regulation of ion transport | 2.41E-05 | | 9.69E-02 | 0.0097 | 0.086 | 24 | 669 | |
| GO:0001508 | action potential | 2.44E-05 | | 9.81E-02 | 0.0097 | 0.086 | 10 | 138 | |
| GO:0055085 | transmembrane transport | 3.00E-05 | | 1.20E-01 | 0.01 | 0.0891 | 38 | 1356 | |
| GO:0042391 | regulation of membrane potential | 3.80E-05 | | 1.53E-01 | 0.0117 | 0.104 | 17 | 391 | |
| GO:0098660 | inorganic ion transmembrane transport | 5.91E-05 | | 2.37E-01 | 0.0169 | 0.15 | 25 | 754 | |
| GO:0098662 | inorganic cation transmembrane transport | 6.56E-05 | | 2.63E-01 | 0.0176 | 0.156 | 23 | 667 | |
| GO:0060081 | membrane hyperpolarization | 7.66E-05 | | 3.07E-01 | 0.0192 | 0.171 | 4 | 17 | |
| GO:0042886 | amide transport | 0.000106 | | 4.24E-01 | 0.0224 | 0.198 | 16 | 384 | |
| GO:0098655 | cation transmembrane transport | 0.000112 | | 4.50E-01 | 0.0225 | 0.2 | 24 | 738 | |
| GO:0098870 | action potential propagation | 0.000143 | | 5.74E-01 | 0.0235 | 0.209 | 3 | 8 | |
| GO:0019227 | neuronal action potential propagation | 0.000143 | | 0.574 | 0.0235 | 0.209 | 3 | 8 | |
| GO:0015672 | monovalent inorganic cation transport | 0.000164 | | 0.658 | 0.0235 | 0.209 | 21 | 617 | |
| GO:0071805 | potassium ion transmembrane transport | 0.000237 | | 0.95 | 0.0272 | 0.241 | 10 | 181 | |
| GO:0071804 | cellular potassium ion transport | 0.000237 | | 0.95 | 0.0272 | 0.241 | 10 | 181 | |
| GO:0007267 | cell-cell signaling | 0.000247 | | 0.991 | 0.0275 | 0.244 | 33 | 1233 | |
| GO:0006814 | sodium ion transport | 0.000257 | | 1 | 0.0278 | 0.247 | 11 | 218 | |
| GO:0006820 | anion transport | 0.000275 | | 1 | 0.029 | 0.257 | 21 | 641 | |
| GO:0035879 | plasma membrane lactate transport | 0.000539 | | 1 | 0.0461 | 0.409 | 3 | 12 | |
| **set2** | **hormone/peptide secrection and regulation** | | | | | | | | |
| GO:0030073 | insulin secretion | 2.32E-05 | | 9.32E-02 | 0.0097 | 0.086 | 13 | 231 | |
| GO:0010817 | regulation of hormone levels | 9.75E-05 | | 3.92E-01 | 0.0224 | 0.198 | 20 | 550 | |
| GO:0050796 | regulation of insulin secretion | 0.000106 | | 4.25E-01 | 0.0224 | 0.198 | 11 | 197 | |
| GO:0015833 | peptide transport | 0.000158 | | 0.636 | 0.0235 | 0.209 | 15 | 357 | |
| GO:0030072 | peptide hormone secretion | 0.000176 | | 0.708 | 0.0244 | 0.217 | 13 | 282 | |
| GO:0090087 | regulation of peptide transport | 0.000224 | | 0.9 | 0.0272 | 0.241 | 13 | 289 | |
| GO:0002790 | peptide secretion | 0.000232 | | 0.931 | 0.0272 | 0.241 | 13 | 290 | |
| GO:0009914 | hormone transport | 0.000233 | | 0.935 | 0.0272 | 0.241 | 15 | 370 | |
| GO:0090276 | regulation of peptide hormone secretion | 0.000485 | | 1 | 0.0461 | 0.409 | 11 | 235 | |
| GO:0046879 | hormone secretion | 0.000522 | | 1 | 0.0461 | 0.409 | 14 | 357 | |
| GO:0046676 | negative regulation of insulin secretion | 0.000535 | | 1 | 0.0461 | 0.409 | 5 | 48 | |
| GO:0002791 | regulation of peptide secretion | 0.000579 | | 1 | 0.0484 | 0.43 | 11 | 240 | |
| **set3** | **muscle contraction and conduction** | | | | | | | | |
| GO:0061337 | cardiac conduction | 2.30E-06 | | 0.00925 | 0.00231 | 0.0205 | 11 | 131 | |
| GO:0003012 | muscle system process | 0.000122 | | 4.89E-01 | 0.0233 | 0.207 | 17 | 430 | |
| GO:0060047 | heart contraction | 0.000143 | | 5.73E-01 | 0.0235 | 0.209 | 13 | 276 | |
| GO:0003015 | heart process | 0.000159 | | 0.638 | 0.0235 | 0.209 | 13 | 279 | |
| GO:0008016 | regulation of heart contraction | 0.000159 | | 0.64 | 0.0235 | 0.209 | 12 | 242 | |
| GO:1903779 | regulation of cardiac conduction | 0.000342 | | 1 | 0.0352 | 0.313 | 6 | 67 | |
| GO:0014888 | striated muscle adaptation | 0.000485 | | 1 | 0.0461 | 0.409 | 5 | 47 | |
| GO:0043501 | skeletal muscle adaptation | 0.000506 | | 1 | 0.0461 | 0.409 | 4 | 27 | |
| **set4** | **others** | | | | | | | | |
| GO:0007626 | locomotory behavior | 2.66E-05 | | 1.07E-01 | 0.0097 | 0.086 | 13 | 234 | |
| GO:0044057 | regulation of system process | 0.000226 | | 0.909 | 0.0272 | 0.241 | 20 | 586 | |
| GO:0007530 | sex determination | 0.000373 | | 1 | 0.0374 | 0.332 | 4 | 25 | |
| GO:0043500 | muscle adaptation | 0.000537 | | 1 | 0.0461 | 0.409 | 7 | 101 | |

**Supplemental Table 5**

| **ID** | **Biological Pathway** | **p-value** | **q-value Bonferroni** | **q-value FDR B&H** | **q-value FDR B&Y** | **Hit in Query List** | **Hit in Genome** |
| --- | --- | --- | --- | --- | --- | --- | --- |
| **set1** |  | | | | | | |
| GO:0019934 | cGMP-mediated signaling | 3.66E-05 | 0.09559 | 0.06971 | 0.5887 | 3 | 13 |
| GO:0071417 | cellular response to organonitrogen compound | 7.12E-05 | 0.186 | 0.06971 | 0.5887 | 11 | 517 |
| GO:0071248 | cellular response to metal ion | 0.0001308 | 0.342 | 0.06971 | 0.5887 | 6 | 151 |
| GO:0071277 | cellular response to calcium ion | 0.00016 | 0.4182 | 0.06971 | 0.5887 | 4 | 53 |
| GO:1901699 | cellular response to nitrogen compound | 0.0002506 | 0.6551 | 0.07279 | 0.6148 | 11 | 597 |
| GO:0071241 | cellular response to inorganic substance | 0.0003002 | 0.7847 | 0.07528 | 0.6358 | 6 | 176 |
| GO:0072503 | cellular divalent inorganic cation homeostasis | 0.0004106 | 1 | 0.08255 | 0.6973 | 9 | 435 |
| GO:0071286 | cellular response to magnesium ion | 0.000543 | 1 | 0.08376 | 0.7075 | 2 | 7 |
| GO:0019935 | cyclic-nucleotide-mediated signaling | 0.0005497 | 1 | 0.08376 | 0.7075 | 4 | 73 |
| GO:0071313 | cellular response to caffeine | 0.0007216 | 1 | 0.08376 | 0.7075 | 2 | 8 |
| GO:0032869 | cellular response to insulin stimulus | 0.0007267 | 1 | 0.08376 | 0.7075 | 6 | 208 |
| GO:0071415 | cellular response to purine-containing compound | 0.0009247 | 1 | 0.08841 | 0.7467 | 2 | 9 |
| GO:0030003 | cellular cation homeostasis | 0.0009471 | 1 | 0.08841 | 0.7467 | 10 | 592 |
| GO:0006873 | cellular ion homeostasis | 0.001144 | 1 | 0.0997 | 0.8421 | 10 | 607 |
| GO:0006874 | cellular calcium ion homeostasis | 0.001284 | 1 | 0.0997 | 0.8421 | 8 | 411 |
| GO:1903792 | negative regulation of anion transport | 0.001317 | 1 | 0.0997 | 0.8421 | 3 | 42 |
| GO:0006875 | cellular metal ion homeostasis | 0.001413 | 1 | 0.0997 | 0.8421 | 9 | 518 |
| GO:0071375 | cellular response to peptide hormone stimulus | 0.001436 | 1 | 0.0997 | 0.8421 | 7 | 324 |
| GO:0055074 | calcium ion homeostasis | 0.001586 | 1 | 0.0997 | 0.8421 | 8 | 425 |
| GO:1901653 | cellular response to peptide | 0.001889 | 1 | 0.1073 | 0.9065 | 7 | 340 |
| GO:0043434 | response to peptide hormone | 0.002027 | 1 | 0.1081 | 0.9132 | 8 | 442 |
| **set2** |  | | | | | | |
| GO:0007411 | axon guidance | 0.0002435 | 0.6365 | 0.07279 | 0.6148 | 7 | 240 |
| GO:0097485 | neuron projection guidance | 0.0002497 | 0.6528 | 0.07279 | 0.6148 | 7 | 241 |
| GO:0061564 | axon development | 0.0003168 | 0.8281 | 0.07528 | 0.6358 | 10 | 514 |
| GO:0016049 | cell growth | 0.0006364 | 1 | 0.08376 | 0.7075 | 10 | 562 |
| GO:0048699 | generation of neurons | 0.0007323 | 1 | 0.08376 | 0.7075 | 18 | 1528 |
| GO:0030182 | neuron differentiation | 0.000737 | 1 | 0.08376 | 0.7075 | 17 | 1398 |
| GO:0007409 | axonogenesis | 0.0007711 | 1 | 0.08399 | 0.7094 | 9 | 475 |
| GO:0031175 | neuron projection development | 0.001192 | 1 | 0.0997 | 0.8421 | 13 | 953 |
| GO:0048812 | neuron projection morphogenesis | 0.00151 | 1 | 0.0997 | 0.8421 | 10 | 630 |
| GO:0022008 | neurogenesis | 0.001515 | 1 | 0.0997 | 0.8421 | 18 | 1628 |
| GO:0060560 | developmental growth involved in morphogenesis | 0.001584 | 1 | 0.0997 | 0.8421 | 6 | 242 |
| GO:0048666 | neuron development | 0.001669 | 1 | 0.0997 | 0.8421 | 14 | 1114 |
| GO:0051960 | regulation of nervous system development | 0.001882 | 1 | 0.1073 | 0.9065 | 12 | 881 |
| **Set 3 (exclude set1 and 2)** | |  | | | | | |
| GO:1902930 | regulation of alcohol biosynthetic process | 0.0001485 | 0.3882 | 0.06971 | 0.5887 | 4 | 52 |
| GO:0003150 | muscular septum morphogenesis | 0.0001567 | 0.4097 | 0.06971 | 0.5887 | 2 | 4 |
| GO:0003149 | membranous septum morphogenesis | 0.0003892 | 1 | 0.08255 | 0.6973 | 2 | 6 |
| GO:0003151 | outflow tract morphogenesis | 0.0004436 | 1 | 0.08282 | 0.6995 | 4 | 69 |
| GO:0060412 | ventricular septum morphogenesis | 0.0005374 | 1 | 0.08376 | 0.7075 | 3 | 31 |
| GO:0072507 | divalent inorganic cation homeostasis | 0.000595 | 1 | 0.08376 | 0.7075 | 9 | 458 |
| GO:0002673 | regulation of acute inflammatory response | 0.0008924 | 1 | 0.08841 | 0.7467 | 4 | 83 |
| GO:0014050 | negative regulation of glutamate secretion | 0.0009247 | 1 | 0.08841 | 0.7467 | 2 | 9 |
| GO:0033084 | regulation of immature T cell proliferation in thymus | 0.001152 | 1 | 0.0997 | 0.8421 | 2 | 10 |
| GO:0002526 | acute inflammatory response | 0.001394 | 1 | 0.0997 | 0.8421 | 5 | 159 |
| GO:0033080 | immature T cell proliferation in thymus | 0.001403 | 1 | 0.0997 | 0.8421 | 2 | 11 |
| GO:0033083 | regulation of immature T cell proliferation | 0.001678 | 1 | 0.0997 | 0.8421 | 2 | 12 |
| GO:0031620 | regulation of fever generation | 0.001678 | 1 | 0.0997 | 0.8421 | 2 | 12 |
| GO:0003205 | cardiac chamber development | 0.001969 | 1 | 0.1076 | 0.9092 | 5 | 172 |
| GO:0033079 | immature T cell proliferation | 0.001977 | 1 | 0.1076 | 0.9092 | 2 | 13 |
| GO:0003279 | cardiac septum development | 0.00213 | 1 | 0.1114 | 0.9405 | 4 | 105 |

**Supplemental Table 6** Correlation between temporal variability of brain regions and age.

|  | **Brain Region** | **R** | **P** |  | **Brain Region** | **R** | **P** |
| --- | --- | --- | --- | --- | --- | --- | --- |
| 1 | HIP.R | -0.2528 | 0.000011 | 46 | PreCG.L | -0.0331 | 0.5704 |
| 2 | PHG.R | -0.2504 | 0.000013 | 47 | IFGoperc.R | -0.0269 | 0.6442 |
| 3 | PoCG.L | -0.1391 | 0.0166 | 48 | HES.R | -0.0255 | 0.6623 |
| 4 | TPOsup.L | -0.1371 | 0.0183 | 49 | SMG.R | -0.0201 | 0.7304 |
| 5 | ORBinf.L | -0.1280 | 0.0276 | 50 | DCG.R | -0.0190 | 0.7442 |
| 6 | TPOmid.L | -0.1251 | 0.0315 | 51 | IFGoperc.L | -0.0184 | 0.7529 |
| 7 | AMYG.R | -0.1238 | 0.0332 | 52 | STG.R | -0.0183 | 0.7532 |
| 8 | IFGtriang.L | -0.1222 | 0.0356 | 53 | ACG.R | -0.0145 | 0.8032 |
| 9 | PreCG.R | -0.1163 | 0.0456 | 54 | IOG.R | -0.0093 | 0.8727 |
| 10 | IPL.R | -0.1150 | 0.0481 | 55 | AMYG.L | -0.0070 | 0.9046 |
| 11 | MFG.L | -0.1150 | 0.0481 | 56 | ORBmid.L | -0.0037 | 0.9492 |
| 12 | PoCG.R | -0.1147 | 0.0486 | 57 | ORBsup.L | -0.0007 | 0.9907 |
| 13 | ANG.L | -0.1134 | 0.0514 | 58 | HES.L | 0.0040 | 0.9457 |
| 14 | SFGmed.L | -0.1119 | 0.0545 | 59 | MTG.R | 0.0050 | 0.9314 |
| 15 | HIP.L | -0.1118 | 0.0547 | 60 | PCG.R | 0.0062 | 0.9157 |
| 16 | PHG.L | -0.1090 | 0.0611 | 61 | SOG.R | 0.0084 | 0.8855 |
| 17 | CAL.R | -0.1048 | 0.0718 | 62 | PCG.L | 0.0097 | 0.8678 |
| 18 | TPOsup.R | -0.1043 | 0.0730 | 63 | IOG.L | 0.0114 | 0.8448 |
| 19 | ORBmid.R | -0.0970 | 0.0958 | 64 | REC.R | 0.0294 | 0.6141 |
| 20 | PCUN.L | -0.0955 | 0.1010 | 65 | ITG.R | 0.0312 | 0.5929 |
| 21 | CAL.L | -0.0936 | 0.1081 | 66 | SOG.L | 0.0334 | 0.5674 |
| 22 | SFGmed.R | -0.0930 | 0.1103 | 67 | ORBsupmed.R | 0.0341 | 0.5589 |
| 23 | CUN.R | -0.0930 | 0.1104 | 68 | REC.L | 0.0391 | 0.5031 |
| 24 | ITG.L | -0.0898 | 0.1234 | 69 | ORBsup.R | 0.0495 | 0.3965 |
| 25 | ORBinf.R | -0.0876 | 0.1326 | 70 | ORBsupmed.L | 0.0570 | 0.3284 |
| 26 | SFGdor.L | -0.0807 | 0.1664 | 71 | INS.R | 0.0616 | 0.2908 |
| 27 | SFGdor.R | -0.0795 | 0.1728 | 72 | INS.L | 0.0618 | 0.2889 |
| 28 | MFG.R | -0.0793 | 0.1736 | 73 | MTG.L | 0.0661 | 0.2573 |
| 29 | LING.R | -0.0781 | 0.1800 | 74 | MOG.R | 0.0672 | 0.2490 |
| 30 | SPG.R | -0.0777 | 0.1824 | 75 | CAU.R | 0.0728 | 0.2119 |
| 31 | IFGtriang.R | -0.0772 | 0.1855 | 76 | SMG.L | 0.0864 | 0.1381 |
| 32 | TPOmid.R | -0.0755 | 0.1954 | 77 | SMA.L | 0.0875 | 0.1331 |
| 33 | PCL.L | -0.0743 | 0.2024 | 78 | ROL.L | 0.0886 | 0.1282 |
| 34 | ACG.L | -0.0740 | 0.2040 | 79 | ROL.R | 0.0964 | 0.0978 |
| 35 | FFG.L | -0.0740 | 0.2044 | 80 | OLF.L | 0.1000 | 0.0859 |
| 36 | PCL.R | -0.0715 | 0.2200 | 81 | SMA.R | 0.1021 | 0.0795 |
| 37 | IPL.L | -0.0669 | 0.2513 | 82 | PAL.R | 0.1030 | 0.0770 |
| 38 | LING.L | -0.0639 | 0.2732 | 83 | THA.R | 0.1034 | 0.0757 |
| 39 | DCG.L | -0.0628 | 0.2813 | 84 | PUT.R | 0.1071 | 0.0657 |
| 40 | PCUN.R | -0.0585 | 0.3161 | 85 | CAU.L | 0.1099 | 0.0589 |
| 41 | ANG.R | -0.0552 | 0.3438 | 86 | OLF.R | 0.1149 | 0.0483 |
| 42 | FFG.R | -0.0530 | 0.3632 | 87 | THA.L | 0.1236 | 0.0335 |
| 43 | STG.L | -0.0525 | 0.3684 | 88 | PAL.L | 0.1246 | 0.0320 |
| 44 | SPG.L | -0.0427 | 0.4646 | 89 | PUT.L | 0.1257 | 0.0306 |
| 45 | CUN.L | -0.0373 | 0.5229 | 90 | MOG.L | 0.1335 | 0.0216 |

**Supplemental Table 7** The 180 areas of the cortical parcellation with index number, short name, description from HCP.

| **Parcel Index** | **Area Name** | **Area Description** | **Parcel Index** | **Area Name** | **Area Description** | **Parcel Index** | **Area Name** | **Area Description** |
| --- | --- | --- | --- | --- | --- | --- | --- | --- |
| **1** | V1 | Primary Visual Cortex | **61** | a24 | Area a24 | **121** | ProS | ProStriate Area |
| **2** | MST | Medial Superior Temporal Area | **62** | d32 | Area dorsal 32 | **122** | PeEc | Perirhinal Ectorhinal Cortex |
| **3** | V6 | Sixth Visual Area | **63** | 8BM | Area 8BM | **123** | STGa | Area STGa |
| **4** | V2 | Second Visual Area | **64** | p32 | Area p32 | **124** | PBelt | ParaBelt Complex |
| **5** | V3 | Third Visual Area | **65** | 10r | Area 10r | **125** | A5 | Auditory 5 Complex |
| **6** | V4 | Fourth Visual Area | **66** | 47m | Area 47m | **126** | PHA1 | ParaHippocampal Area 1 |
| **7** | V8 | Eighth Visual Area | **67** | 8Av | Area 8Av | **127** | PHA3 | ParaHippocampal Area 3 |
| **8** | 4 | Primary Motor Cortex | **68** | 8Ad | Area 8Ad | **128** | STSda | Area STSd anterior |
| **9** | 3b | Primary Sensory Cortex | **69** | 9m | Area 9 Middle | **129** | STSdp | Area STSd posterior |
| **10** | FEF | Frontal Eye Fields | **70** | 8BL | Area 8B Lateral | **130** | STSvp | Area STSv posterior |
| **11** | PEF | Premotor Eye Field | **71** | 9p | Area 9 Posterior | **131** | TGd | Area TG dorsal |
| **12** | 55b | Area 55b | **72** | 10d | Area 10d | **132** | TE1a | Area TE1 anterior |
| **13** | V3A | Area V3A | **73** | 8C | Area 8C | **133** | TE1p | Area TE1 posterior |
| **14** | RSC | RetroSplenial Complex | **74** | 44 | Area 44 | **134** | TE2a | Area TE2 anterior |
| **15** | POS2 | Parieto-Occipital Sulcus Area 2 | **75** | 45 | Area 45 | **135** | TF | Area TF |
| **16** | V7 | Seventh Visual Area | **76** | 47l | Area 47l (47 lateral) | **136** | TE2p | Area TE2 posterior |
| **17** | IPS1 | IntraParietal Sulcus Area 1 | **77** | a47r | Area anterior 47r | **137** | PHT | Area PHT |
| **18** | FFC | Fusiform Face Complex | **78** | 6r | Rostral Area 6 | **138** | PH | Area PH |
| **19** | V3B | Area V3B | **79** | IFJa | Area IFJa | **139** | TPOJ1 | Area TemporoParietoOcci pital Junction 1 |
| **20** | LO1 | Area Lateral Occipita 1 | **80** | IFJp | Area IFJp | **140** | TPOJ2 | Area TemporoParietoOcci pital Junction 2 |
| **21** | LO2 | Area Lateral Occipital 2 | **81** | IFSp | Area IFSp | **141** | TPOJ3 | AreaTemporoParietoOccipital Junction 3 |
| **22** | PIT | Posterior InferoTemporal | **82** | IFSa | Area IFSa | **142** | DVT | Dorsal Transitional Visual Area |
| **23** | MT | Middle Temporal Area | **83** | p9- 46v | Area posterior 9-46v | **143** | PGp | Area PGp |
| **24** | A1 | Primary Auditory Cortex | **84** | 46 | Area 46 | **144** | IP2 | Area IntraParietal 2 |
| **25** | PSL | PeriSylvian Language Area | **85** | a9- 46v | Area anterior 9-46v | **145** | IP1 | Area IntraParietal 1 |
| **26** | SFL | Superior Frontal Language Area | **86** | 9-46d | Area 9-46d | **146** | IP0 | Area IntraParietal 0 |
| **27** | PCV | PreCuneus Visual Area | **87** | 9a | Area 9 anterior | **147** | Pfop | Area PF opercular N |
| **28** | STV | Superior Temporal Visual Area | **88** | 10v | Area 10v | **148** | PF | Area PF Complex |
| **29** | 7Pm | Medial Area 7P | **89** | a10p | Area anterior 10p | **149** | PFm | Area PFm Complex |
| **30** | 7m | Area 7m | **90** | 10pp | Polar 10p | **150** | PGi | Area PGi |
| **31** | POS1 | Parieto-Occipital Sulcus Area 1 | **91** | 11l | Area 11l | **151** | PGs | Area PGs |
| **32** | 23d | Area 23d | **92** | 13l | Area 13l | **152** | V6A | Area V6A |
| **33** | v23ab | Area ventral 23 a+b | **93** | OFC | Orbital Frontal Complex | **153** | VMV1 | VentroMedial Visual Area 1 |
| **34** | d23ab | Area dorsal 23 a+b | **94** | 47s | Area 47s | **154** | VMV3 | VentroMedial Visual Area 3 |
| **35** | 31pv | Area 31p ventral | **95** | LIPd | Area Lateral IntraParietal dorsal | **155** | PHA2 | ParaHippocampal Area 2 |
| **36** | 5m | Area 5m | **96** | 6a | Area 6 anterior | **156** | V4t | Area V4t |
| **37** | 5mv | Area 5m ventral | **97** | i6-8 | Inferior 6-8 Transitional Area | **157** | FST | Area FST |
| **38** | 23c | Area 23c | **98** | s6-8 | Superior 6-8 Transitional Area | **158** | V3CD | Area V3CD |
| **39** | 5L | Area 5L | **99** | 43 | Area 43 | **159** | LO3 | Area Lateral Occipital 3 |
| **40** | 24dd | Dorsal Area 24d | **100** | OP4 | Area OP4/PV | **160** | VMV2 | VentroMedial Visual Area 2 |
| **41** | 24dv | Ventral Area 24d | **101** | OP1 | Area OP1/SII | **161** | 31pd | Area 31pd |
| **42** | 7AL | Lateral Area 7A | **102** | OP2-3 | Area OP2-3/VS | **162** | 31a | Area 31a |
| **43** | SCEF | Supplementary and Cingulate Eye Field | **103** | 52 | Area 52 | **163** | VVC | Ventral Visual  Complex |
| **44** | 6ma | Area 6m anterior | **104** | RI | RetroInsular Cortex | **164** | 25 | Area 25 |
| **45** | 7Am | Medial Area 7A | **105** | PFcm | Area PFcm | **165** | s32 | Area s32 |
| **46** | 7Pl | Lateral Area 7P | **106** | PoI2 | Posterior Insular Area 2 | **166** | pOFC | posterior OFC Complex |
| **47** | 7PC | Area 7PC | **107** | TA2 | Area TA2 | **167** | PoI1 | Area Posterior Insular 1 |
| **48** | LIPv | Area Lateral IntraParietal ventral | **108** | FOP4 | Frontal OPercular Area 4 | **168** | Ig | Insular Granular Complex |
| **49** | VIP | Ventral IntraParietal Complex | **109** | MI | Middle Insular Area | **169** | FOP5 | Area Frontal Opercular 5 |
| **50** | MIP | Medial IntraParietal Area | **110** | Pir | Pirform Cortex | **170** | p10p | Area posterior 10p |
| **51** | 1 | Area 1 | **111** | AVI | Anterior Ventral Insular Area | **171** | p47r | Area posterior 47r |
| **52** | 2 | Area 2 | **112** | AAIC | Anterior Agranular Insula Complex | **172** | TGv | Area TG Ventral |
| **53** | 3a | Area 3a | **113** | FOP1 | Frontal OPercular Area 1 | **173** | MBelt | Medial Belt Complex |
| **54** | 6d | Dorsal area 6 | **114** | FOP3 | Frontal OPercular Area 3 | **174** | LBelt | Lateral Belt Complex |
| **55** | 6mp | Area 6mp | **115** | FOP2 | Frontal Opercular Area 2 | **175** | A4 | Auditory 4 Complex |
| **56** | 6v | Ventral Area 6 | **116** | PFt | Area PFt | **176** | STSva | Area STSv anterior |
| **57** | p24pr | Area Posterior 24 prime | **117** | AIP | Anterior IntraParietal Area | **177** | TE1m | Area TE1 Middle |
| **58** | 33pr | Area 33 prime | **118** | EC | Entorhinal Cortex | **178** | PI | Para-Insular Area |
| **59** | a24pr | Anterior 24 prime | **119** | PreS | PreSubiculum | **179** | a32pr | Area anterior 32 prime |
| **60** | p32pr | Area p32 prime | **120** | H | Hippocampus | **180** | p24 | Area posterior 24 |
